# Type 1 diabetes risk genes mediate pancreatic beta cell survival in response to proinflammatory cytokines

**DOI:** 10.1101/2021.10.29.466025

**Authors:** Paola Benaglio, Han Zhu, Mei-Lin Okino, Jian Yan, Ruth Elgamal, Naoki Nariai, Elisha Beebe, Katha Korgaonkar, Yunjiang Qiu, Margaret Donovan, Joshua Chiou, Jacklyn Newsome, Jaspreet Kaur, Sierra Corban, Anthony Aylward, Jussi Taipale, Bing Ren, Kelly A Frazer, Maike Sander, Kyle J Gaulton

## Abstract

Beta cells intrinsically contribute to the pathogenesis of type 1 diabetes (T1D), but the genes and molecular processes that mediate beta cell survival in T1D remain largely unknown. We combined high throughput functional genomics and human genetics to identify T1D risk loci regulating genes affecting beta cell survival in response to the proinflammatory cytokines IL-1β, IFNγ, and TNFα. We mapped 38,931 cytokine-responsive candidate *cis*-regulatory elements (cCREs) active in beta cells using ATAC-seq and single nuclear ATAC-seq (snATAC-seq), and linked cytokine-responsive beta cell cCREs to putative target genes using single cell co-accessibility and HiChIP. We performed a genome-wide pooled CRISPR loss-of-function screen in EndoC-βH1 cells, which identified 867 genes affecting cytokine-induced beta cell loss. Genes that promoted beta cell survival and had up-regulated expression in cytokine exposure were specifically enriched at T1D loci, and these genes were preferentially involved in inhibiting inflammatory response, ubiquitin-mediated proteolysis, mitophagy and autophagy. We identified 2,229 variants in cytokine-responsive beta cell cCREs altering transcription factor (TF) binding using high-throughput SNP-SELEX, and variants altering binding of TF families regulating stress, inflammation and apoptosis were broadly enriched for T1D association. Finally, through integration with genetic fine mapping, we annotated T1D loci regulating beta cell survival in cytokine exposure. At the 16p13 locus, a T1D variant affected TF binding in a cytokine-induced beta cell cCRE that physically interacted with the *SOCS1* promoter, and increased *SOCS1* activity promoted beta cell survival in cytokine exposure. Together our findings reveal processes and genes acting in beta cells during cytokine exposure that intrinsically modulate risk of T1D.

## INTRODUCTION

Type 1 diabetes (T1D) is a complex disease characterized by autoimmune destruction of the insulin-producing beta cells in the pancreas. During progression to T1D, immune infiltration and inflammation occurs in the local environment around beta cells, through which beta cells are directly exposed to external stimuli such as proinflammatory cytokines secreted by immune cells^1^. Beta cells themselves intrinsically contribute to the development of T1D in response to these stimuli. Studying beta cell function during T1D progression directly is challenging due to the limited availability of samples and the difficulty in capturing the precise window in which beta cells are the target of immune attack. An alternate strategy is to model T1D progression *in vitro*, for example by culturing islets or beta cells with pro-inflammatory cytokines interleukin 1β (IL-1β), interferon γ (IFNγ), and tumor necrosis factor α (TNFα)^2–6^. Application of this model has revealed widespread effects on beta cell gene regulation, function and survival in response to cytokine exposure^2,4–7^. However, the genes and processes in beta cells that directly contribute to the development of T1D in the context of cytokine exposure remain poorly defined.

Human genetics represents an avenue through which to identify genes and processes within beta cells that play a causal role in T1D. Genome-wide association studies have identified over 90 genomic regions associated with T1D, the majority of which are non-coding and likely affect gene regulation^8,9^. Variants at T1D risk loci are enriched in islet *cis-*regulatory elements (cCREs) induced by pro-inflammatory cytokine exposure6, but not islet regulatory elements in the basal state, which supports that risk of T1D in beta cells acts downstream of external stimuli during disease progression. Genes at several T1D risk loci have been shown to affect beta cell function in cytokine signaling such *PTPN2* and *DEXI*^6,10,11^. At most T1D loci, however, whether risk genes mediate beta cell function in cytokine exposure is unknown. More broadly, determining the pathways through which these risk genes operate can help to converge on mechanisms through which beta cells intrinsically affect disease.

In this study we used a suite of functional genomics assays to map *cis*-regulatory programs in pancreatic beta cells as well as identify genes that affect beta cell survival upon exposure to the pro-inflammatory cytokines IL1β, IFNγ and TNFα. We then integrated these data with fine-mapping data to identify T1D risk variants regulating beta cell survival during cytokine exposure.

## RESULTS

### Overview of study design

In this study we combined human genetics and functional genomics to identify genes that affect risk of T1D by modulating pancreatic beta cell survival in response to proinflammatory cytokine exposure (**Figure 1**). First, we created a map of cytokine-responsive *cis*-regulatory elements (cCREs) in pancreatic beta cells treatment using bulk and single nuclear ATAC-seq. Second, we linked cytokine-responsive beta cell cCREs to target genes using single cell co-accessibility and HiChIP. Third, we identified genes affecting beta cell survival in cytokine exposure using a genome-wide CRISPR knockout screen in EndoC-βH1 cells. Fourth, we identified functional variants in cytokine-responsive beta cell chromatin by assaying *in vitro* transcription factor binding using high-throughput SNP-SELEX. Finally, we integrated these functional genomics data with fine-mapping of 136 T1D signals to annotate functional T1D risk variants directly regulating genes involved in cytokine-induced beta cell survival.

**Figure 1.**
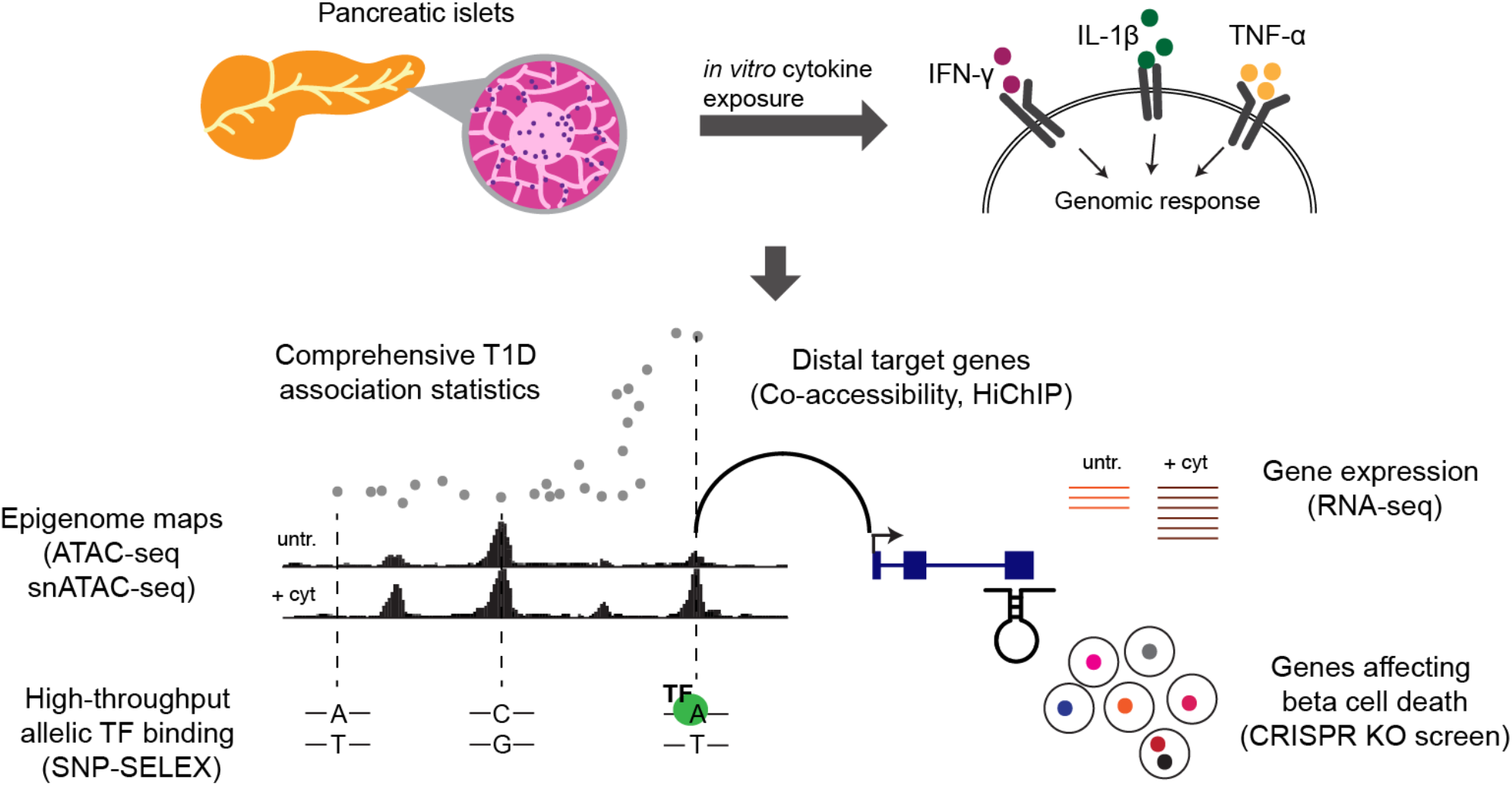
Overview of study design. Schematic representation of the experimental design to model inflammation of human pancreatic islet and characterization of changes using multiple genome-wide functional assays to identify mechanisms involved in type 1 diabetes risk.

### Map of pancreatic beta cell chromatin in response to cytokines

To identify epigenomic changes in pancreatic islets in response to cytokine exposure, we performed ATAC-seq in a total of 7 primary islet preparations cultured *in vitro* with the cytokines IL-1β, IFNγ, and TNFα as well as in untreated conditions (**Supplementary Table 1**). We performed these assays across multiple dimensions of cytokine treatment (35 assays in total), including different treatment doses (high-dose: 0.5 ng/mL IL-1β, 10 ng/mL IFN-γ, 1 ng/mL TNF-α; low-dose: 0.01 ng/mL IL-1β, 0.2 ng/mL IFN-γ, and 0.02 ng/mL TNF-α), duration (6hr, 24hr, 48hr, 72hr), and cytokines used (3 cytokines: IL-1β, IFN-γ and TNFα, or 2 cytokines: IL-1β and IFN-γ).

We determined the effects of inflammatory cytokine signaling for all treatments on islet accessible chromatin genome-wide by performing principal component analysis (PCA) using normalized read counts (**Figure 2a**). There were reproducible patterns of chromatin accessibility across replicate samples of the same treatment, with clear separation between the cytokine-treated and untreated samples. We also observed patterns across different cytokine treatments, where the low-dose cytokine had an intermediate effect to the high-dose cytokine treatment. For example, at the *CXCL10/11* locus there was a notable gradient of increasing accessible chromatin signal across untreated, low-dose cytokine and high-dose cytokine treated samples (**Figure 2b**).

**Figure 2.**
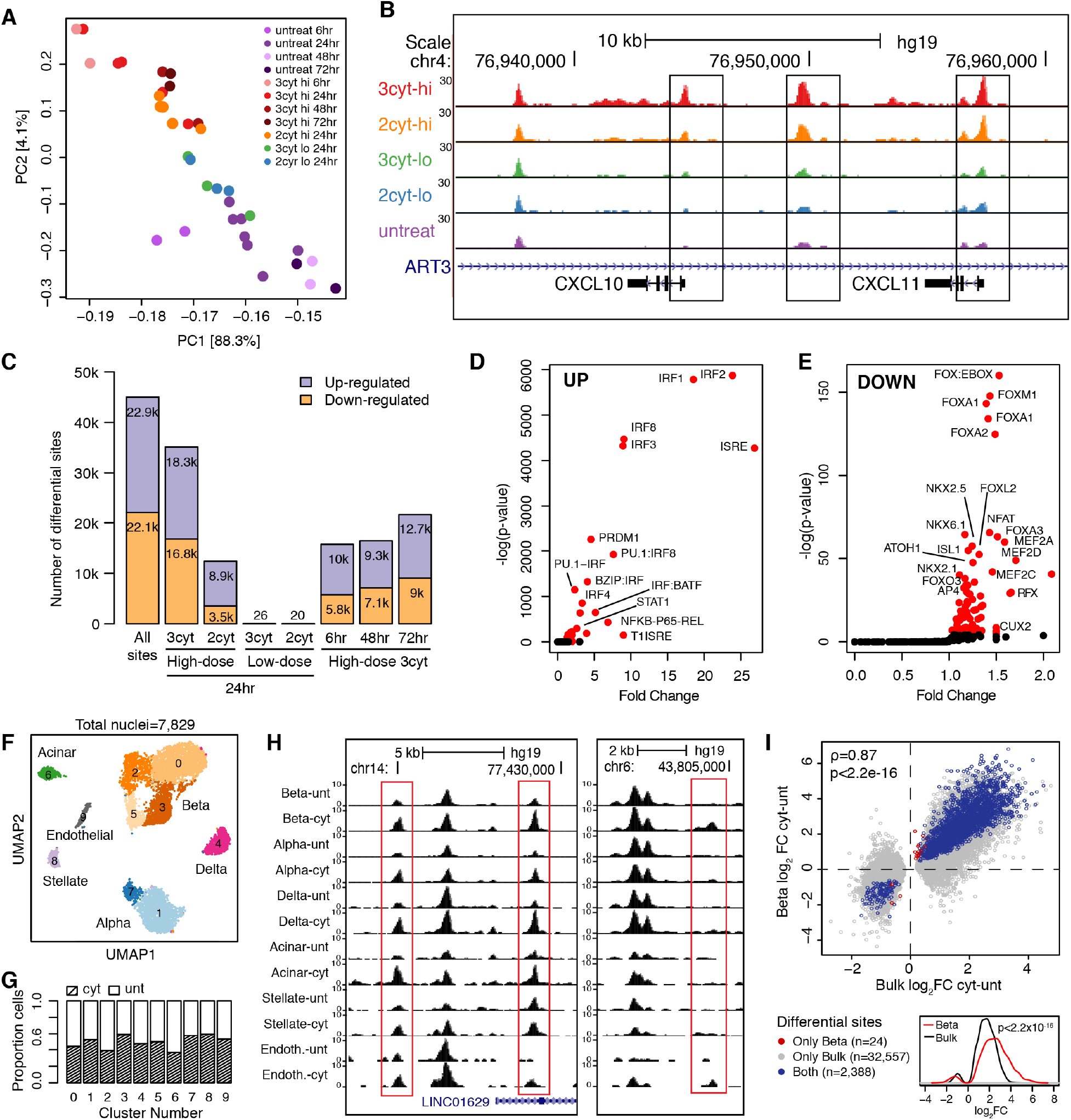
Map of islet accessible chromatin upon exposure to inflammatory cytokines. A) Principal component analysis showing distribution of samples (n=35) based on the different cytokine treatments, color-coded as shown in the legend. Hi: high dose; Lo: low dose; 3cyt: IL-1β, IFN-γ, TNFa; 2cyt: IL-1β, IFN-γ. B) Genome-browser screenshot at the *CXCL10*/*CXCL11* locus, showing ATAC-seq tracks combined across cytokine treatments at 24hrs. The example shows increased chromatin accessibility at cCRE with higher doses and number of cytokines used for stimulation. C) Number of differential cCRE for each treatment compared to control, and union of all differential sites. D-E) Sequence TF motifs enriched in all up-regulated (D) and all down-regulated (E) cCRE compared to all cCRE. F) Clustering of single cell accessible chromatin profiles of islet samples from 4 individuals. Cells are plotted based on the first two UMAP components. G) Barplot showing the proportion of cytokine treated and untreated cells in each cluster. H) Genome-browser screenshots showing example of cytokine-induced cCREs with constitutive effect in all cell types (left) or specific effect in beta cells (right). I) Scatterplot showing effect of cytokine-resposive cCREs in bulk ATAC (x-axis) and in beta cell snATAC (y-axis). Spearman correlation coefficient and p-values are indicated. Bottom: density plot showing increased effect size in beta cells, across DACs significant in both snATAC for beta cells and bulk ATAC. Wilcoxon signed rank test p-value is shown.

We next identified islet cCREs with significant differences in chromatin accessibility in cytokine-treated compared to untreated cells. We first defined a set of 165,884 cCREs genome-wide active in islets. From these 165,884 cCREs, we next identified cCREs with differential accessibility in cytokine treatment compared to control using DESeq2^12^. There were 22,877 cCREs with increased activity in any cytokine treatment and 22,092 cCREs with decreased activity in any cytokine treatment (FDR<0.1, **Figure 2c, Supplementary Table 2**). Notably, there was a marked difference in the number of cytokine-responsive cCREs across treatment dose, with almost no such cCREs at low-dose treatment (**Figure 2c**). When comparing treatments with and without TNFα including TNFα produced a broadly stronger effect on cytokine-responsive cCREs overall (**Supplementary Figure 1a**), although there were no cCREs with significant changes in activity between the two conditions, suggesting modest effects on individual cCREs. Finally, we identified 1,000 cCREs with differential activity across duration (**Supplementary Figure 1b**, p-value <0.01, linear regression), the majority with increased accessibility with longer duration of treatment (**Supplementary Figure 1c**).

In order to identify transcriptional regulators of cytokine-responsive cCREs in islets, we performed sequence motif enrichment using HOMER^13^. Consistent with previous reports^6^, cCREs with increased activity in cytokine treatment were strongly enriched for IRF (IRF1 *P*<10^−300^, IRF2 *P*<10^−300^), STAT (STAT1 *P*=2.8×10^−130^) and NFkB (NFKB-P65-REL *P*=2.1×10^−279^) motifs (**Figure 2d**, **Supplementary Table 3**). Conversely, cCREs with decreased activity in any cytokine treatment were most enriched for FOXA (*P*=5.8×10^−63^), NKX6.1 (*P*=1.1×10^−28^), NFAT (*P*=3.8×10^−29^), and MEF2 (*P*=9.2×10^−27^), motifs (**Figure 2e**, **Supplementary Table 3**). We next identified sequence motifs with variable enrichment across different dimensions of cytokine treatment. For example, motifs with variable enrichment in up-regulated cCREs across duration of cytokine treatment included SMAD family TFs, which had stronger enrichment at later timepoints (SMAD2 6hr *P* =0.24, 24hr *P* =0.03, 48hr *P* =8.6×10^−4^, 72hr *P* =2.2×10^−6^), and motifs with variable enrichment among down regulated cCREs included RFX, NFAT and MEF2 family motifs (**Supplementary Figure 1d**).

The effects of cytokine exposure on individual islet cell types are obscured from assays of bulk tissue. Therefore, we next performed single nuclear ATAC-seq (snATAC-seq) in cytokine-treated and untreated islets from four donors at 24 hours post-treatment. We used high-dose of all three cytokines IL-1β, IFNγ, and TNFα as the treatment for these assays, as this produced the strongest effects in bulk assays. After extensive quality control, which included removal of low quality and doublet cells (see **Methods**), we performed UMAP dimensionality reduction and clustering on a total of 7,829 nuclei, which identified 9 clusters (**Figure 2f**). Each cluster contained cells from all four donors and was equally represented by untreated (total nuclei = 3,947) and cytokine-treated (total nuclei = 3,882) cells (**Figure 2g, Supplementary Figure 2a-b**). We assigned each cluster cell type identity based on accessibility levels at the promoters of known marker genes (**Supplementary Figure 2c-d**), which revealed endocrine alpha, beta and delta cells as well as exocrine, endothelial and stellate cells.

We next defined cCREs in beta and other cell types and used the resulting cCREs to annotate the cytokine-responsive cCREs identified in bulk ATAC-seq (**Supplementary Figure 3a**). We identified 38,931 cytokine-responsive islet cCREs active in beta cells, a small percentage (8.2%) of which were specific to beta cells relative to other endocrine cell types (example in **Figure 2h**). We further used snATAC data from cytokine-treated and untreated cells to identify differential sites in beta cells directly. There were 2,412 cytokine-responsive beta cell cCREs (FDR<0.1 **Supplementary Figure 3b, Supplementary Table 2**), almost all of which (99%, 2,388) had significant and concordant effects in bulk islets. The effects of cytokine treatment on cCRE activity were generally stronger on beta cells relative to bulk islets, although there were fewer cCREs overall with significant changes in activity in beta cells likely owing to the smaller number of samples (**Figure 2i**). Compared to alpha cells, there were substantially more cCREs with cytokine-responsive activity in beta cells (2,412 vs. 226), despite having similar total numbers of cells (**Supplementary Figure 3c**). Furthermore, the effects of cytokine treatment on cCRE activity were consistently stronger in beta cells compared to alpha cells (Wilcox signed rank test P=1.2×10^−255^) (**Supplementary Figure 3d**). These results suggest that beta cell chromatin is more responsive to pro-inflammatory stress than chromatin in other islet cell types.

**Figure 3.**
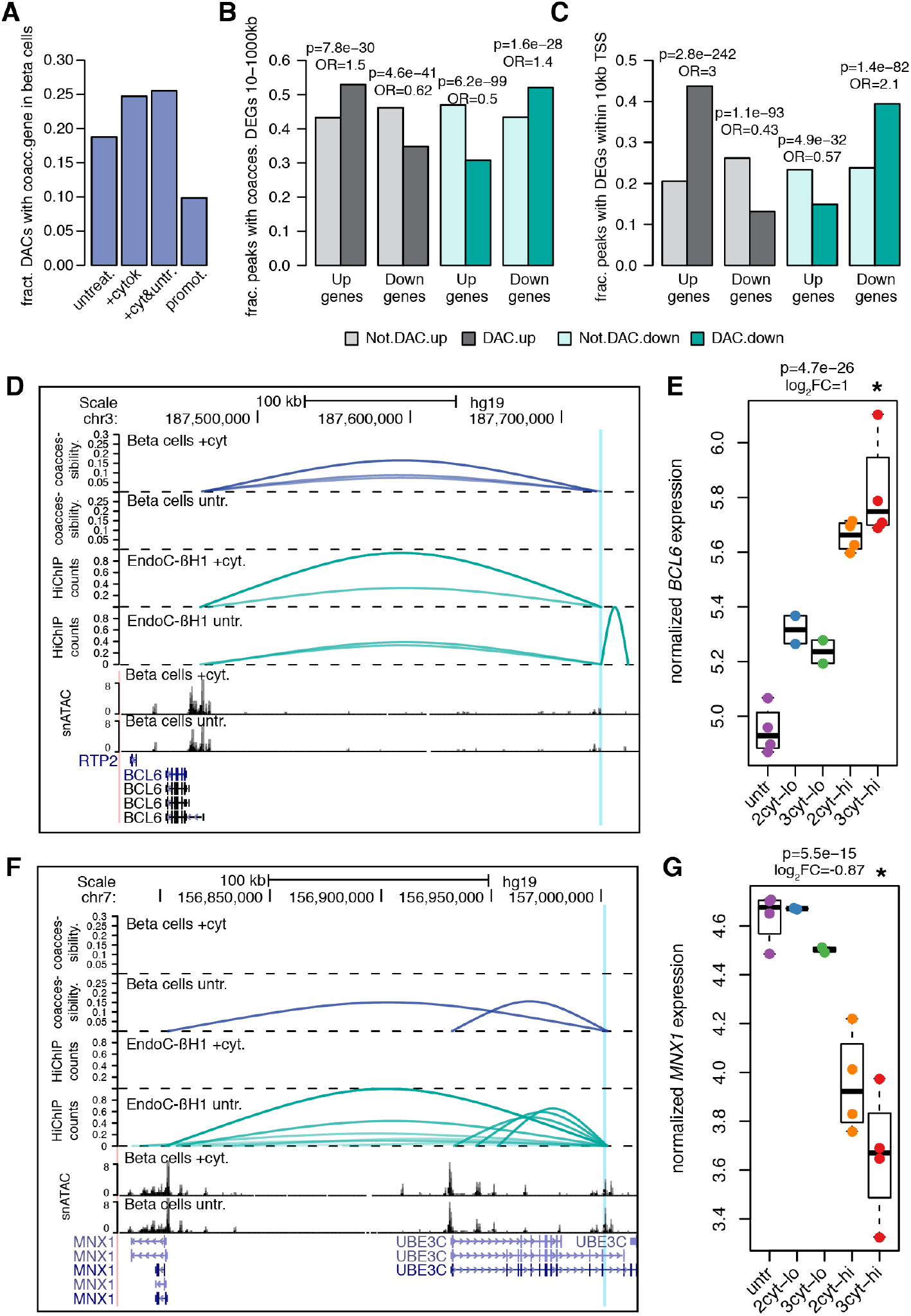
Target genes of beta cells cCREs in inflammatory cytokine exposure. A) Fraction of cytokine-responsive cCREs (Differentially Accessible cCREs-DACs) either co-accessible with at least one gene promoter in beta cells in untreated, cytokine-stimulated or pooled conditions; or proximal to a promoter (<10kb TSS). B) Enrichment of distal DACs (>10kb from TSS) for co-accessibility to genes with concordant cytokine-induced effects. Fisher’s exact test p-values and odds ratios are shown. Co-accessibility was calculated from pooled cytokine-treated and untreated beta cells. C) Enrichment of promoter-proximal DACs (<10kb from TSS) for genes with concordant cytokine-induced effects. Fisher’s exact test p-values and odds ratios are shown. D-E) Example of a cytokine up-regulated peak (blue vertical line) with HiChIP-validated co-accessibility with the promoter of a cytokine-upregulated gene (*BCL6*). D) From top to bottom: co-accessibility in beta cells in cytokine or untreated conditions, virtual 4C profiles from HiChIP in EndoC-βH1 in cytokine or untreated conditions, snATAC profiles in beta cells in cytokine or untreated conditions, gene annotations. Only co-accessibility arcs that link the highlighted distal peak and a promoter peak are shown. E) *Voom*-normalized and batch-corrected expression of *BCL6* in human islet samples after different cytokine treatment conditions. DESeq p-value and log_2_ fold change are from DESeq differential expression between 3-cytokine, high-doses treated islets (red) vs untreated (purple). F-G) Same as D-E, but showing an example of a cytokine down-regulated peak (blue vertical line) with HiChIP-validated co-accessibility with the promoter of a cytokine-downregulated gene (*MNX1*). 2cyt: cytokine treatment with IL1b and IFNg, 3cyt: cytokine treatment with IL1b, IFNg and TNFa, lo: low-dose, hi: high-dose, untr: untreated

Finally, we identified TF motifs differentially enriched in cytokine-responsive beta cell accessible chromatin. We identified motifs differentially enriched in single cytokine-treated and untreated beta cells using ChromVAR^14^. The most enriched motifs in beta cells were broadly consistent with those identified in bulk data, with IRF-family TFs showing highest enrichment in cytokine-treated beta cells and FOXA TFs the strongest depletion (**Supplementary Figure 3e, Supplementary Table 3**). However, when comparing motif enrichments in alpha and beta cells there was more significant enrichment of IRF- and STAT-family motifs in cytokine-treated beta cells, further supporting that cytokine treatment has stronger effects in beta cells (**Supplementary Figure 3e**).

In summary, we generated a comprehensive catalog of cCREs that respond to pro-inflammatory cytokines in pancreatic islets and beta cells.

### Linking cytokine-responsive beta cell cCREs to target genes

As the majority of cytokine-responsive beta cell cCREs are distal to gene promoters, we next sought to link cytokine-responsive cCREs to the target genes they regulate in beta cells.

We first identified cytokine-responsive cCREs correlated with the activity of gene promoter cCREs using co-accessibility across single cytokine-treated and untreated beta cells with Cicero^15^. In total, we identified 400,403 and 277,447 pairs of co-accessible cCREs (score >0.05), in cytokine-treated and untreated beta cells, respectively, 30% of which involved a gene promoter cCRE. We then annotated cytokine-responsive beta cell cCREs co-accessible with at least one gene promoter. There were 11,124 and 8,434 cytokine-responsive cCREs co-accessible with a putative target gene in cytokine-treated and untreated beta cells, respectively, while ~10% of cytokine-responsive cCREs were at promoters directly (**Figure 3a**). As co-accessibility represents a correlation between cCREs that may not always reflect direct *cis* regulation, we next mapped 3D physical interactions between cCREs using HiChIP in high-dose cytokine-treated (IL1β, IFNγ and TNFα) and untreated EndoC-βH1 beta cells. Co-accessible sites were significantly enriched for 3D interactions compared to non-co-accessible sites (Fisher’s exact test P<2.2×10^−16^; cytokine treated OR=3.6; untreated OR=3.2). In total, 2,520 and 2,063 distal cCREs co-accessible with a promoter in cytokine-treated and untreated cells, respectively, had a 3D interaction (FDR<.10).

We next assessed the relationship between the activity of cytokine-responsive beta cell cCREs and the expression of target genes linked to the cCREs in cytokine treatment. We performed RNA-seq in islets treated with high- and low-dose cytokines for 24hr and identified differentially expressed genes (DEGs) in cytokine-treated compared to untreated cells. High-dose exposure to all three cytokines (IL1β, IFNγ and TNFα) produced the largest changes in expression, where 3,367 genes had increased, and 3,414 genes had decreased expression in cytokine-treated compared to untreated islets (**Supplementary Figure 4a-f, Supplementary Table 4**). High-dose treatment using just IL1β and IFNγ resulted in 5,051 differentially expressed genes. As with bulk ATAC-seq data, low-dose treatment resulted in fewer differentially expressed genes overall (330 with three cytokines, 324 with two cytokines), and these genes were largely a subset of the genes identified in high dose treatment (**Supplementary Figure 4b-c, Supplementary Table 4**).

We determined whether genes co-accessible with cytokine-response distal cCREs had directionally concordant changes in expression. Here we used just genes differentially expressed in high-dose cytokine treatment. Distal cCREs (>10kb from TSS) with up-regulated or down-regulated activity in cytokine treatment were significantly enriched for co-accessibility to genes with increased or decreased expression, respectively (**Figure 3b**). We observed similar patterns when considering distal cCREs with 3D physical interactions to genes (**Supplementary Figure 4g**). Cytokine-responsive cCREs proximal to gene promoters were also enriched for concordant effects on expression with stronger enrichment than for distal cCREs (**Figure 3c**). At the 3q27 locus, a cytokine-induced beta cell cCRE was co-accessible with the *BCL6* promoter in cytokine-treated beta cells, and *BCL6* attenuates the proinflammatory response but induces apoptosis in beta cells^16^ (**Figure 3d**). The beta cell cCRE interacted with the *BCL6* promoter in cytokine-treated cells only (cytokine-treated FDR=6.2×10^−6^), and *BCL6* had increased expression in cytokine treatment (**Figure 3d-e**). Similarly, at the 7q36 locus, a beta cell cCRE with decreased activity in cytokine exposure was co-accessible with the promoter of *MNX1,* which is involved in maintaining beta cell fate (**Figure 3f**). We observed an interaction between the beta cell cCRE and the *MNX1* promoter in untreated beta cells only (untreated FDR=5.4×10^−6^) and *MNX1* had decreased expression in cytokine treatment (**Figure 3g-f).**

Together these results reveal the target genes of cytokine-responsive distal cCRE activity in beta cells.

### Genes affecting beta cell survival in response to cytokine exposure

Given target genes of cytokine-responsive cCREs in beta cells, we next determined which genes had cellular functions directly relevant to T1D pathogenesis. As beta cell loss is the primary pathogenic endpoint of T1D, we sought to identify genes affecting beta cell survival in response to cytokine exposure. We therefore performed a genome-wide pooled CRISPR loss-of-function screen in the human EndoC-βH1 beta cell line using cell survival under cytokine exposure (high-dose IL-1β, IFN-γ, TNF-α) for 72hr as an endpoint. We selected a longer duration of treatment than for chromatin and gene expression assays to effectively capture cell loss in response to cytokine treatment. In brief, after transfecting cells with the GeCKO v2 CRISPR single guide RNA (sgRNA) library, we split and cultured cells in either high-dose cytokine or control (0.1% BSA). The representation of sgRNAs between cytokine-treated and untreated conditions was compared to identify genes promoting or preventing beta cell loss in response to cytokine exposure. An overview of the screen design is shown in **Figure 4a**.

**Figure 4.**
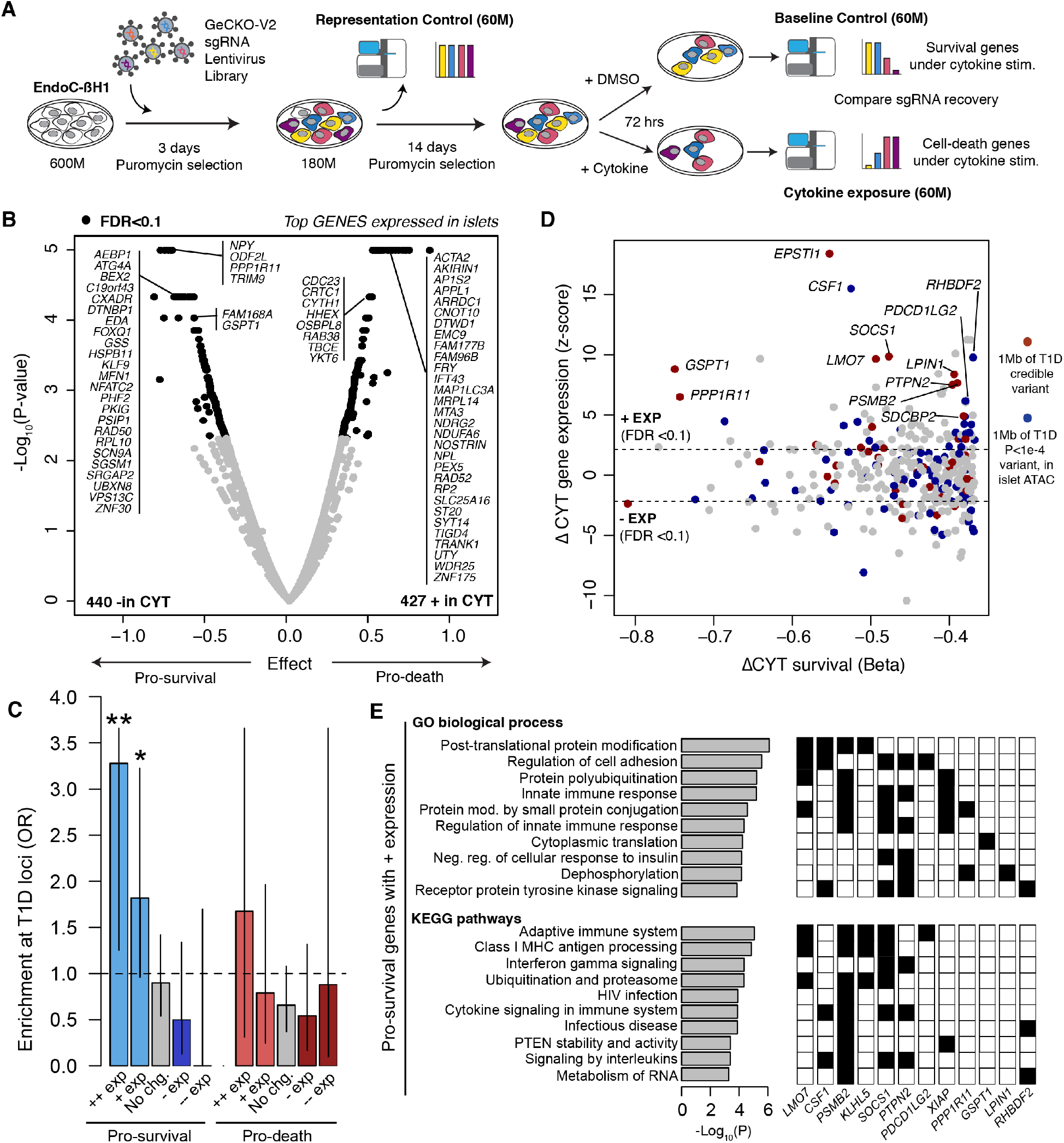
Genes affecting beta cell survival in cytokine exposure. A) Overview of the genome-wide CRISPR loss-of-function screen in cytokine-treated EndoC-βH1 cells. B) Volcano plot of genes with significant (FDR<0.1) enrichment and depletion from the screen. Labeled genes include the top genes with an average TPM expression >1 in islets. C) Enrichment of known T1D risk loci for genes enriched and depleted in screen, partitioned by differential expression (+/− exp = FDR<0.1, ++/−-exp = FDR<0.1 and *P*<1×10^-5^) in islets after cytokine stimulation (3-cyt, high-doses). Values are odds ratio and error bars are 95% confidence interval from Fisher’s exact test. D) Scatterplot showing the effect size (beta) of genes promoting beta survival in the screen and differential expression of the gene in islets after cytokine treatment (z-scores). Genes mapping within 1MB of a known T1D locus or within 1MB of a variant with nominal (p<1×10-4) T1D association are colored. E) Molecular pathways from gene ontology (GO) and KEGG enriched in genes with increased expression in cytokine-treated islets and promoting beta cell survival. A subset of genes mapping to known T1D loci or loci with nominal T1D association are shown with corresponding pathway annotations. Only pathways that contain at least one T1D gene are shown, while the full list is shown in Supplementary Table 6.

Among 18,703 genes targeted by sgRNAs (6 sgRNAs per gene) after transduction, 867 genes had significant (FDR<.10) differences in recovered sgRNAs between cytokine-treated and untreated cells. Among these, sgRNAs for 427 genes were enriched in cytokine-treated compared to untreated cells and therefore these genes promoted beta cell loss (‘pro-death’) in response to cytokine exposure (**Figure 4b**, **Supplementary Table 5**). Conversely, sgRNAs for 440 genes were depleted in cytokine-treated compared to untreated cells and therefore these genes prevented beta cell loss (‘pro-survival’) in response to cytokine exposure (**Figure 4b**, **Supplementary Table 5**). The results of our screen identified genes previously shown to affect survival in beta cells, for example *XIAP*^17^, *JUND*^18^*, PTPN2*^10^, and *SOCS1*^19,20^. To annotate the function of pro-death and pro-survival genes, we performed gene ontology enrichment analyses (**Supplementary Table 6**). As expected, pro-death genes were enriched for terms related to DNA damage response, apoptosis and protein folding, and pro-survival genes were enriched for autophagy, which protects against beta cell stress, and phosphorylation and kinase activity, which suppress inflammatory responses. Pro-survival genes were also enriched for RNA metabolism and splicing and pro-death genes were enriched for lipid metabolism, processes which have all been implicated in beta cell function and survival^21,22^.

Interestingly, genes regulating processes related to mitochondrial function were highly enriched among both pro-death and pro-survival genes. We found that pro-survival mitochondria-related genes were primarily involved in mitochondria organization and mitophagy, such as *USP36, VDAC1, MFF, TIMM9, YME1L1, SIRT5,* and *SPATA18*. Conversely, mitochondria-related genes in the pro-death category were mostly electron transport chain components, such as *NDUFA6*, *NDUFB2*, *ACAD9*, *CYCS* and *SDHD*. A key mitophagy regulator, *CLEC16A*, has been previously shown to protect beta cells against inflammatory damage, mediated in part by reactive oxygen species (ROS) generated in beta cell mitochondria^23^. Our data suggest that mitophagy and mitochondria quality control are important pro-survival processes in beta cells in response to pro-inflammatory cytokines and provide novel regulators of beta cell mitophagy.

Given genes and molecular processes affecting cytokine-induced beta cell loss, we next determined which genes and processes might be relevant to T1D pathogenesis. First, we tested for enrichment of genes affecting cytokine-induced beta cell loss at loci involved in genetic risk of T1D. We observed no evidence for enrichment among the full set of either pro-survival or pro-death genes. Next, we further segregated pro-survival and pro-death genes based on whether their expression was significantly up-regulated or down-regulated, or had no change, in cytokine exposure. Pro-survival genes that had up-regulated expression in cytokines (n=84 genes) were significantly enriched at known T1D loci (+exp OR=1.82, 95% CI=0.97,3.23 *P*=.048, Fisher’s test), and no other subset showed any enrichment (**Figure 4c**). This enrichment was stronger when considering only genes with the largest increases (P<1×10^−5^) in cytokine-induced expression (++exp OR=3.28, 95% CI=1.33,7.37 *P=*5.1×10^−3^). Numerous genes with highly induced expression mapped to known T1D risk loci such as *PTPN2, EPSTI1, SOCS1, PSMB2, PPP1R11, LPIN1* and *LMO7* (**Figure 4d**). This subset of genes also included several with roles in mitophagy such as *NBR1* and *MFN1*.

We next characterized the molecular functions of the pro-survival genes with up-regulated expression in cytokine exposure. These genes were broadly enriched for molecular processes related to modulation of the inflammatory response, ubiquitination and proteasomal degradation, translation, and autophagy (**Supplementary Table 6**, **Figure 4e**). Among genes at T1D loci were negative regulators of cytokine signaling *PTPN2* and *SOCS1*, both of which function by inhibiting JAK/STAT signaling to suppress inflammatory responses and promote beta cell survival^10^. Other beta cell survival genes such as *KLHL5, LMO7, NEDD4L, ASB2* and *PPP1R11* function in protein ubiquitination, which targets proteins for degradation by the proteasome, and *PSMD2* is a component of the 20S proteasome itself. Proinflammatory cytokines induce endoplasmic reticulum (ER) stress in beta cells^24^, and proteasome-mediated ER-associated protein degradation (ERAD) resolves ER stress in beta cells^25^. Ubiquitin-mediated proteolysis may therefore protect beta cells from cytokine-induced stress, although the function of most of these genes in beta cells is unknown. We also observed enrichment of class I MHC antigen-related terms, although genes annotated with these terms were largely overlapping with other terms.

Together these results identify genes and molecular processes that affect beta cell loss in response to proinflammatory cytokine exposure and reveal that T1D risk is specifically enriched for pro-survival genes highly induced in cytokine exposure.

### Identifying functional regulatory variants in beta cell cCREs with SNP-SELEX

Given that beta cell pro-survival genes up-regulated in cytokine exposure were enriched at T1D risk loci, we next sought to determine the transcriptional regulators of gene activity in beta cells during cytokine exposure through which T1D genetic risk is mediated.

Because risk variants often affect transcriptional regulation via differential transcription factor binding (TF), we systematically determined the effects of genetic variants in cytokine-responsive beta cell cCREs on TF binding. A total of 184,086 variants were tested for *in vitro* differential TF binding using a highly multiplexed assay SNP-SELEX^26^. The 184,086 variants were selected based on mapping in islet enhancer regions genome-wide (56,796) or mapping to known diabetes risk loci (T1D: 86,067, T2D: 33,354), in addition to variants randomly selected across the genome (7,869). Among the tested variants were 183,373 SNPs and 713 indels (insertions/deletions of 1-3 bp). We designed a library of 44bp oligos surrounding each variant containing each of the four possible alleles for SNPs, or each of the two observed alleles for indels. We then tested oligos for binding to 530 distinct *E. coli*-expressed TF proteins by sequencing recovered oligos across four binding cycles, where the entire experiment was performed in duplicate (**Figure 5a**; **Supplementary Table 7**).

**Figure 5.**
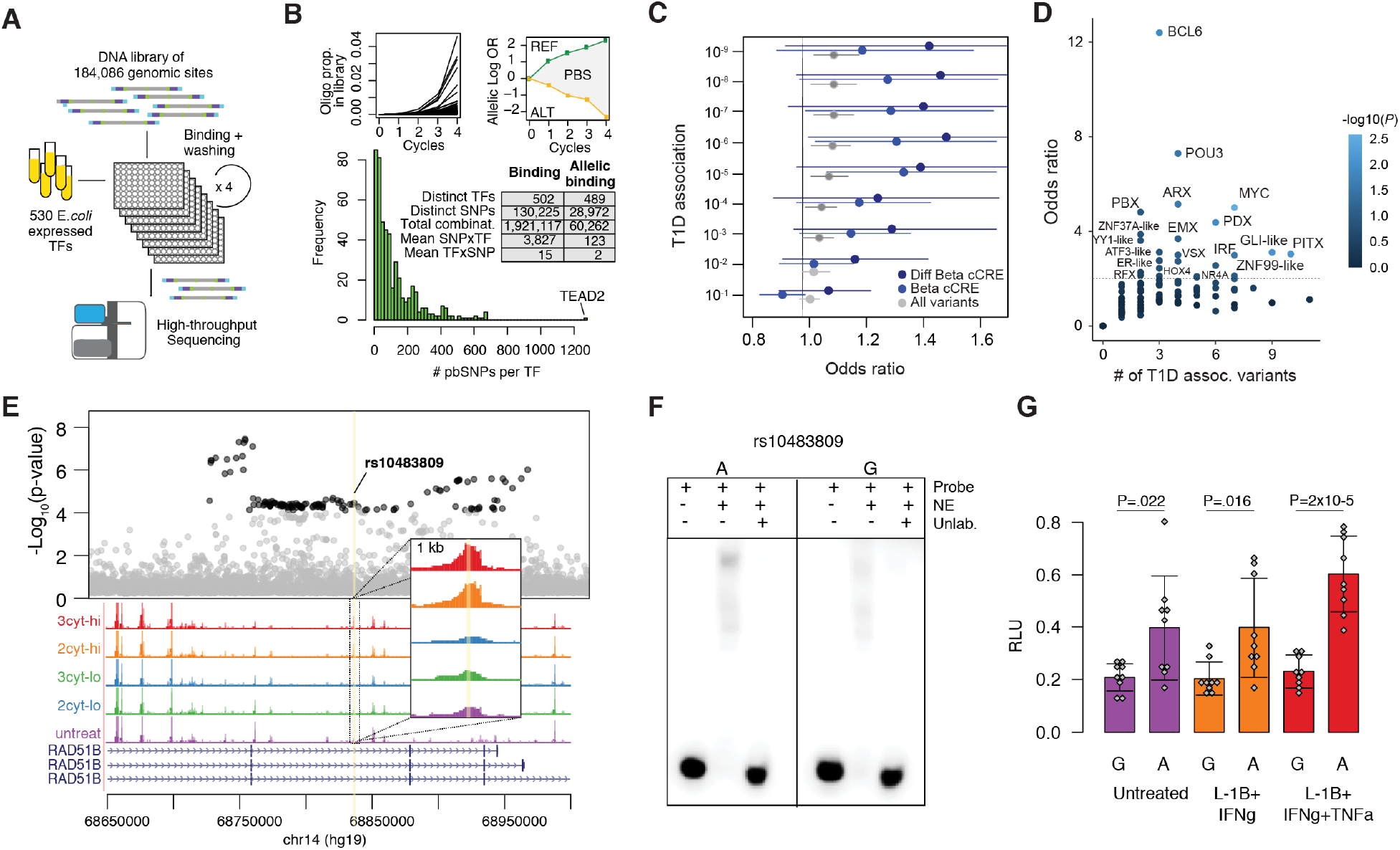
Identifying transcriptional regulators affecting T1D risk in beta cell cCREs with SNP-SELEX. A) Overview of the HT-SELEX–Seq experiment. B) Top: Example of enrichment profiles of bound oligos within an experiment and of one SNP with preferential binding to one nucleotide. Bottom: Distribution of the number of variants with allelic binding per TF across 489 TFs and table summarizing the number of bound variants and significant allelic binding variants across TFs. C) Enrichment of variants with allelic binding for T1D association among all tested variants, variants in beta cell cCREs and variants in cytokine-induced beta cell cCREs. Values represent odds ratio and 95% CI by Fisher’s exact test. D) Enrichment of variants with allelic binding of specific TF sub-families for T1D association among variants in cytokine-induced beta cell cCREs. Values represent odds ratio by Fisher’s exact test, and points are colored by p-value. E) Regional plot of association p-values with T1D, with variants with p-value <10^−4^ in black; bulk ATAC-seq tracks from human islets 2cyt: cytokine treatment with IL1b and IFNg, 3cyt: cytokine treatment with IL1b, IFNg and TNFa, lo: low-dose, hi: high-dose, untr: untreated. A zoom in to the candidate variant is shown in the panel. F) Electrophoretic mobility shift assay using nuclear extract (NE) from MIN6 cells with probes for the alleles of rs10483809. G) Luciferase assays for rs10483809 in untreated MIN6 cells or after 24h treatment with high-dose cytokines. Relative light units (RLUs) are normalized to cells transfected with the empty vector (pGL4.23). Average and standard deviation of 6 transfection replicates are shown. *P*-values from two-tailed *t*-tests are shown.

After applying quality-filtering criteria (**Supplementary Figure 5a-c**), a total of 130,225 variants were bound by at least one TF and were further analyzed for differential allelic binding. We identified variants with allelic differences in TF binding from SNP-SELEX by calculating a preferential binding score (PBS) score between alleles (see Methods) (**Figure 5b**, **Methods**). There were 28,972 variants that affected binding of at least one TF (P<0.05 by Monte Carlo randomization), with a mean of 2 TFs per variant and of 123 variants per TF (**Figure 5b**). TFs from the same family often clustered together based on the correlation in variant effects on binding (**Supplementary Figure 5d**)^26–28^. Variant effects on TF binding from SNP-SELEX were generally correlated with predicted effects including from DeepSEA^29^ (mean r=0.81, **Supplementary Figure 5e**) and position weight matrix (PWM) models (r=0.91, **Supplementary Figure 5f**), although this was also highly variable across TFs (**Supplementary Figure 5g**). Consistent with previous findings^26^, a minority of variants (29% on average per TF) with allelic effects from SNP-SELEX had a corresponding PWM prediction (**Supplementary Figure 5h**), which highlights the benefit of this experimentally generated resource.

In total, there were 8,424 variants in beta cell cCREs affecting TF binding, including 2,229 in cytokine-responsive beta cell cCREs. We determined if variants affecting TF binding within beta cell cCREs were enriched for T1D association at different p-value thresholds compared to other tested variants. T1D-associated variants in beta cell cCREs were enriched for allelic effects on TF binding, and this enrichment was stronger for variants in cytokine-responsive cCREs (**Figure 5c**). By comparison, there was limited enrichment among all tested variants for allelic effects on TF binding (**Figure 5c**). We next grouped TFs into 220 sub-families using TFClass^30^, and tested for enrichment of T1D association among variants in cytokine-responsive beta cell cCREs disrupting TF binding in each sub-family. TF sub-families with strongest enrichment (OR>2) included BCL6, POU3, PBX, MYC, ARX and PDX1 (**Figure 5d**). We also observed enrichment for sub-families regulating stress, mitophagy and immune responses such as ATF3-like, IRF, NR4, and GLI-like TFs (**Figure 5d**). To identify specific TFs likely regulating cytokine-induced beta cell cCREs, we annotated TF genes in each sub-family with differential expression in cytokine exposure. TF genes within enriched sub-families with cytokine-induced expression included *BCL6*, *GLIS3*, *IRF1/2/*7/*9*, *PBX1, PDX1, ATF3, NR4A1/3,* and *MYC* (**Supplementary Table 4**).

We then identified specific variants at T1D risk loci affecting TF binding in cytokine-responsive beta cell cCREs. In total 380 variants in cytokine-responsive beta cell cCREs mapped within 1MB of a known T1D locus and affected TF binding, including for TF with differential expression in cytokine stimulation. For example, at the *RAD51B* locus, variant rs10483809 (T1D P=8.1×10^−6^) mapped in a cytokine-induced beta cell cCRE and the T1D risk allele had preferential binding to IRF- and CUX-family TFs (**Figure 5e**). As SNP-SELEX is based on *in vitro* interactions, we validated allelic effects on regulatory activity in beta cells directly. Electrophoretic mobility shift assay (EMSA) demonstrated protein binding to the T1D risk allele using nuclear extract from the beta cell line MIN6 (**Figure 5f**). We also identified increased enhancer activity for the risk allele in luciferase gene reporter assays in MIN6 cells, which was more pronounced in cytokine stimulation (**Figure 5g**). This variant maps in *RAD51B* which is a pro-apoptotic protein involved in DNA recombination^31^ and up-regulated in cytokine-treated islets (**Supplementary Table 4**), although did not affect cell death in our screen.

Together these results identify functional variants altering TF binding in beta cell cCREs and reveal transcriptional regulators through which variants in cytokine-responsive beta cell cCREs broadly affect T1D risk.

### T1D risk variants linked to genes affecting beta cell survival in cytokines

Finally, given the molecular processes and regulatory networks enriched for T1D risk in cytokine-induced beta cells, we layered functional genomics together with human genetics data to annotate specific T1D loci that regulate genes affecting beta cell loss in cytokine exposure.

We first intersected cytokine-responsive beta cell cCREs with fine-mapping 99% credible sets of 136 T1D signals^8^. At 77 T1D signals, at least one credible set variant overlapped a beta cell cCRE, and at 52 signals a credible set variant overlapped a cytokine-responsive beta cell cCRE (**Supplementary Table 8**). Among T1D signals with credible set variants in cytokine-responsive beta cell cCREs were those at loci previously implicated in beta cell function such as *PTPN2*, *DEXI/CLEC16A, GLIS3* and *DLK1*^10,11,32–34^. For the T1D signals with credible set variants in cytokine-induced beta cell cCREs, we next linked variants at 37 signals to putative target genes using beta cell co-accessibility (**Supplementary Table 8**). Genes linked to credible set variants in cytokine-responsive beta cell cCREs included 19 genes affecting beta cell loss from the CRISPR screen in addition to several key stress response genes (**Supplementary Table 8**).

At the *DEXI/CLEC16A* (16p13) locus, which has two independent T1D risk signals, seven credible set T1D variants from the secondary signal overlapped cytokine-induced beta cell cCREs (**Figure 6a-c, Supplementary Table 8**). Among these, only one variant rs35342456 had significant allelic effects on TF binding from SNP-SELEX (P=2.4×10^−5^), and therefore is a functional candidate for underlying the association signal at this locus (**Figure 6d**). A previous study identified a functional variant rs193778 in cytokine-stimulated islet chromatin at this locus^6^, but this variant was not present in our 99% credible set data. To validate that rs35342456 has regulatory effects in beta cells, we performed an EMSA to measure TF binding to each allele using nuclear extract from cytokine-treated and untreated MIN6 beta cells (**Figure 6e, Supplementary Figure 6**). Consistent with the SNP-SELEX data, we observed allele-specific effects of this variant on TF binding in beta cells (**Figure 6e**).

**Figure 6.**
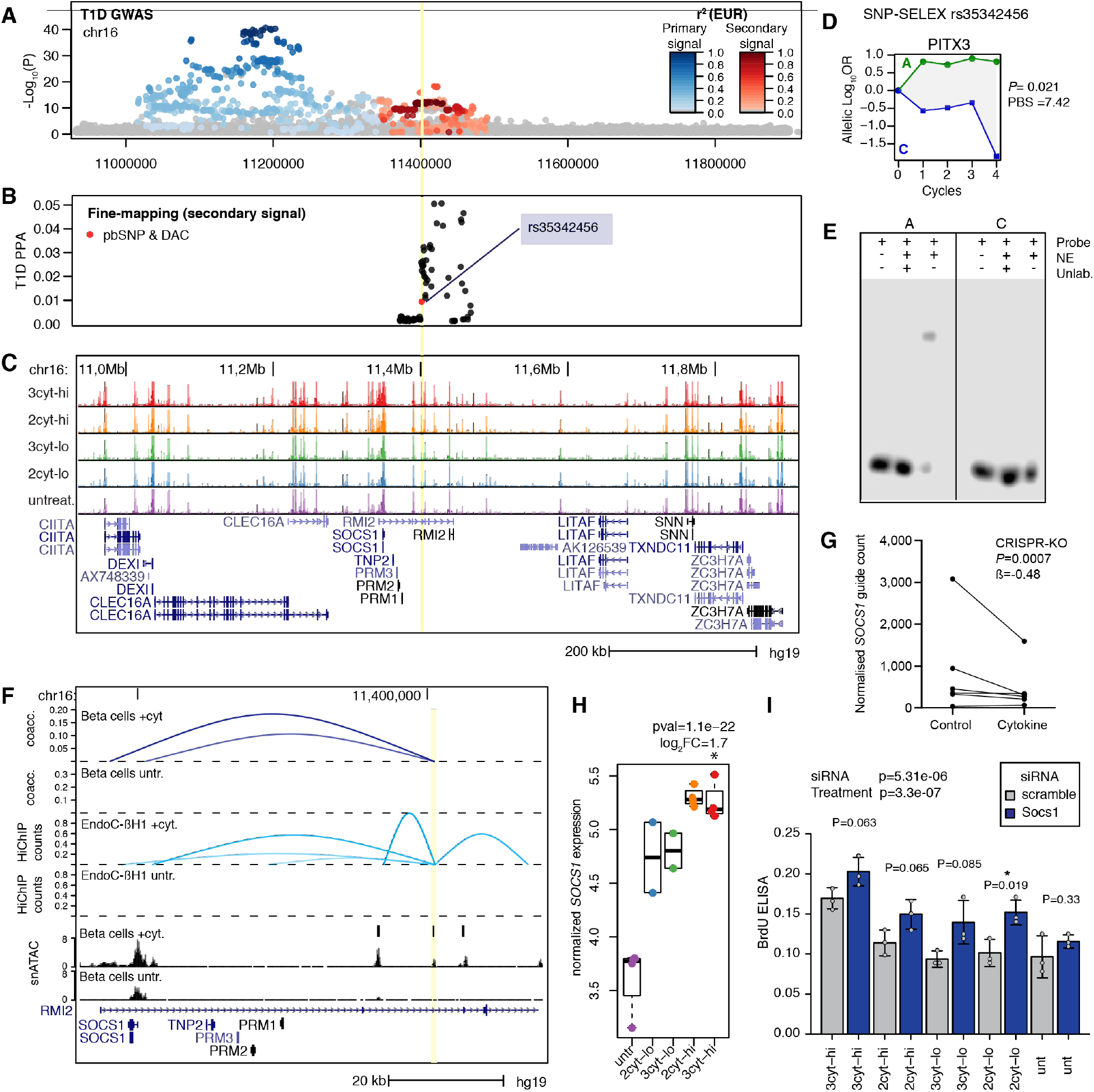
T1D locus 16p13 regulates beta cell survival gene *SOCS1* in cytokine exposure. A) T1D GWAS regional association plot showing two independent signals at the *DEXI/SOCS1* locus. B) Fine mapping posterior probabilities of the secondary signal. Variants with SNP-SELEX significant effect on differential TF binding and within a cytokine-responsive cCRE are highlighted in red. C) Genome-browser of the *DEXI/SOCS1* locus showing bullk ATAC-seq tracks from human islets after the indicated treatments and gene annotations. D) SNP-SELEX results for the candidate variant rs35342456. 2cyt: cytokine treatment with IL1b and IFNg, 3cyt: cytokine treatment with IL1b, IFNg and TNFa, lo: low-dose, hi: high-dose, untr: untreated. E) EMSA with nuclear extract (NE) from MIN6 cells showing preferential binding to probes with the reference allele, consistent with SNP-SELEX results. F) Zoom in of the locus showing location of the candidate variant rs35342456 (yellow vertical line) in a cytokine-up regulated beta cell peak that is co-accessible and show HiChIP interaction with *SOCS1* promoter in cytokine-treated beta cells and EndoC-βH1 respectively. G) Count number of each sgRNA in the CRISPR-KO screen targeting *SOCS1* in untreated ang high-cytokine treated Endoβ-CH1, normalised to the sequencing depth of each sample, showing higher counts in untreated samples for five out of six sgRNAs. Counts for the same sgRNA in the two conditions are connected with a line. H) Normalized and batch-corrected expression of *SOCS1* in human islet samples after different cytokine treatment conditions. DESeq p-value and log_2_ fold change are from DESeq, showing significant (FDR<0.1) higher expression between 3-cytokine, high-doses treated islets (red) vs untreated (purple). H) Effect of *Socs1* knock-down by siRNA on cell death-induced DNA fragmentation using an anti-BrdU ELISA in MIN6 cells. ANOVA p-values testing effect of both siRNA and treatment are shown on the top. Two-tailed t-test p-values are shown for pairs of Socs1/scramble siRNA in each treatment condition. Error bars represent standard deviation. 2cyt: cytokine treatment with IL1b and IFNg, 3cyt: cytokine treatment with IL1b, IFNg and TNFa, lo: low-dose, hi: high-dose, untr: untreated.

The cytokine-induced cCRE harboring rs35342456 was co-accessible with the promoter region of *SOCS1*, which had up-regulated expression and promoted beta cell survival in cytokine exposure, implicating *SOCS1* as a candidate causal gene for T1D risk (**Figure 6f-h**). We confirmed *SOCS1* as a target of cytokine-dependent cCRE activity at this locus using the HiChIP data in cytokine-treated and untreated EndoC-βH1 beta cells. We observed a significant interaction (FDR=0.068) between the cCRE and *SOCS1* promoter in cytokine-treated beta cells, which is highlighted using virtual 4C centered on the cCRE (**Figure 6f**). By comparison, there was no evidence for an interaction between the cCRE and *SOCS1* promoter in untreated beta cells (**Figure 6f**). Furthermore, there was no evidence of interaction between the cCRE and the promoter regions of other genes at the locus, including previously implicated candidate genes *DEXI* and *CLEC16A,* the expression of which was also not significantly affected by cytokine treatment (**Supplementary Table 4**). These results reveal that *SOCS1* is a likely *cis-*regulatory target of T1D risk variant activity in cytokine-induced beta cells at the 16p13 locus.

In the CRISPR screen *SOCS1* promoted beta cell survival after cytokine exposure, and *SOCS1* had significant increase in cytokine-induced expression (**Figure 6g-h**). We determined the effects of *SOCS1* on cytokine-induced beta cell survival using an independent assay that measures cell death via DNA fragmentation (**see Methods**). We performed siRNA-mediated knockdown of *Socs1* in the pancreatic beta cell line MIN6 cultured with high-dose or low-dose cytokine treatment with or without TNFα as well as in untreated conditions, and measured DNA fragmentation using an anti-BrdU ELISA. We observed significant increase in DNA fragmentation in *Socs1* siRNA compared to scramble control siRNA (two-way ANOVA P=3.3×10^−7^, **Figure 6i**). Furthermore, the effects of *Socs1* knockdown were more pronounced in cytokine-treated compared to untreated beta cells, although this interaction was not significant (P>.05) (**Figure 6i**).

These results reveal that the up-regulation of *SOCS1* activity in response to cytokine exposure promotes beta cell survival and plays a likely causal role in risk of T1D.

## DISCUSSION

Combining functional genomics and genetic association data revealed genes and molecular processes involved in risk of T1D in beta cells. Genes with highly induced expression that promoted beta cell survival in response to cytokine exposure were specifically enriched at T1D loci. These genes broadly reflect two classes of intrinsic mechanisms that protect beta cells against proinflammatory cytokines: first, direct inhibition of the inflammatory response and, second, resolution of stress-induced damage due to the inflammatory response. The activity of these pro-survival genes is induced by distal beta cell cCREs that respond to cytokine signaling, and these cCREs in turn often harbor T1D risk variants. As a result, risk of T1D can likely be explained in part by reduced induction of pro-survival genes in beta cells in response to proinflammatory cytokines during disease progression.

Pro-survival genes involved in modulating the immune response included *PTPN2* and *SOCS1,* which map to known T1D risk loci. Both *PTPN2* and *SOCS1* suppress the inflammatory response by inhibiting the JAK/STAT pathway. Previous studies in model systems demonstrated that knockdown of *PTPN2* in beta cells led to increased phosphorylation of STAT1/3 upon activation by interferon gamma as well as phosphorylation of the pro-apoptotic protein BIM^10,35^, which in turn increased beta cell death. Studies in model systems have shown that *SOCS1* promotes beta cell survival by blocking the phosphorylation of JAK to suppress the inflammatory response^19,20,36^. In line with these findings in model organisms, our study reveals a role for *PTPN2* and *SOCS1* in promoting cell survival in human beta cells in cytokine exposure. Furthermore, based on direct links to a T1D risk variant, the regulation of *SOCS1* activity in beta cells after cytokine exposure likely plays a causal role in T1D.

Pro-survival genes with highly induced expression in cytokines were also involved ubiquitin-mediated proteolysis. Among these were several *LMO7, PPP1R11* and *PSMD2* which mapped to T1D loci^36–38^. Cytokine signaling in beta cells induces proteasomal activity^39^, and the proteasome is involved in cell survival^39–42^. As proinflammatory cytokines induce ER stress in beta cells in the context of T1D^2^ which can lead to cell death^43^, and ER stress is resolved in in part through protein degradation^44^, these genes may function in resolving ER stress. However, at present, the function of these genes in beta cells and the mechanisms through which ubiquitination-mediated proteolysis may regulate survival in T1D are unclear. In addition to ER stress, cytokines also cause beta cell death through the production of ROS in mitochondria^45–47^. Mitophagy is induced by ROS production downstream of inflammation to prevent beta cell damage^48^, and our analyses revealed pro-survival genes affecting mitophagy. Moreover, many pro-survival genes were also involved in class I MHC antigen processing and presentation. Class I MHC activity in beta cells is necessary for progression of T1D^49^, likely by an increased exposure of beta cell antigens promoting an immune response. It is thus possible that some genes play dual roles in T1D via beta cell survival as well as antigen presentation, although other models will be needed to test the latter hypothesis.

While our study identifies *SOCS1* as a novel candidate gene for T1D, several other genes at the 16p13 locus have been previously implicated in disease including *DEXI* and *CLEC16A*. Inhibition of *DEXI* in beta cells was previously shown to reduce the activation of STAT and chemokine production and promote survival in response to viral double-stranded RNA (dsRNA)^11^. In our CRISPR screen we observed the opposite effect where *DEXI* function promoted beta cell survival. T1D risk variants at 16p13 were also previously shown to physically interact with the *DEXI* promoter in cytokine-treated beta cells, although we did not find corresponding evidence in our HiChIP data in this study, nor did *DEXI* have differential expression in cytokine exposure. In the case of *CLEC16A,* pancreas-specific deletion in mice led to decreased mitophagy and abnormal mitochondria^33^, although we didn’t identify *CLEC16A* in our screen and *CLEC16A* expression was not affected in cytokine treatment. Another candidate gene at this locus, *CIITA,* is an MHC class II *trans*-activator that has induced expression in cytokine treatment and is expressed in beta cells from T1D donors^50^. Therefore, it is likely that several genes mediate T1D risk in beta cells at this locus.

Multiple genes such as *DEXI* had opposite effects on beta cell survival compared to previous reports. *DEXI* is a pro-survival gene in our screen but was previously shown to induce beta cell death in response to viral dsRNA^11^. In another example, *NDRG2* is a pro-death gene in our data but was previously shown to protect beta cells from lipotoxicity^51^. In such cases, opposing effects on survival could arise from differences in the cellular responses to different stressors such as viral dsRNA, cytokines, or lipids, or from differences between human and mouse. In addition, our screen identified genes affecting beta cell proliferation, such as *NFATC2*^52^. The relevance of such genes to primary beta cell function in cytokine exposure is unclear, however, as these genes might reflect the transformed nature of EndoC-βH1 cells. At present, EndoC-βH1 is the only human beta cell option for a genome wide CRISPR screen, which require large numbers of cells for sufficient coverage. Study designs that compare sgRNAs recovered from cell populations FACS-sorted based on a cellular marker, as was recently done for insulin content in EndoC-βH1 cells^53^, may also complement the cell loss design used here. Human pluripotent stem cell (hPSC) derived islet organoids could be a future platform for screens but will require a differentiation-compatible lentivirus transduction method and scalable beta cell purification strategy.

Our study also provided novel insight into transcriptional regulators of gene activity in beta cells through which T1D risk variants act in response to cytokine signaling. Variants with differential transcription factor binding within cytokine-responsive beta cells cCREs were broadly enriched for T1D association, supporting that a subset of T1D risk acts in beta cells by disrupting transcriptional programs which respond to cytokine signaling. The function of TF sub-families most enriched for T1D association largely mirrored the molecular processes enriched in beta cell pro-survival genes with induced expression, providing orthogonal support for the role of these processes in T1D risk in beta cells. Furthermore, by annotating TF genes within these sub-families with altered expression in cytokine exposure, we pinpointed specific TFs likely driving beta cell *cis*-regulatory activity affecting T1D risk. For example, in beta cells, IRF TFs and *BCL6* regulate inflammation^16,54,55^, *ATF3*, *GLIS3*, and *MYC* regulate stress response and apoptosis^56–59^, and *NR4A1*, *NR4A3, PDX1* and *MYC* regulate mitochondrial function and mitophagy^58,60,61^. These findings also support a model in which T1D risk variants broadly affect the activity of a wide variety of TFs regulating disease-relevant pathways in beta cells.

While IL1β, IFNγ and TNFα have been extensively used as an *in vitro* model of T1D, beta cells are exposed to additional cytokines during disease progression. For example, a recent study revealed widespread changes in beta cell regulatory programs after exposure to IFNα^62^, which is involved in anti-viral immunity. Continued generation of genomic maps in beta cells exposed to other disease-relevant cytokines will therefore be informative in interpreting T1D risk. Beta cells are also exposed to other T1D-relevant external stimuli beyond cytokines. *In vitro* models of endoplasmic reticulum stress^63^, oxidative stress^64^, hypoxia^65^ and hyperglycemia^66^ have all been used to study beta cell function in the context of disease but the genomic response of beta cells to these stressors and their role in T1D genetic risk remains largely unknown. As *in vitro* models only partially re-capitulate disease biology, mapping regulatory programs in beta cells from individuals in the early stages of T1D will also help in interpreting disease risk.

In summary we identified transcriptional regulators, genes and molecular pathways that affect T1D risk by modulating beta cell survival after cytokine exposure, providing new avenues to preserve beta cell mass in T1D.

## METHODS

### Human islet samples

Human islet samples obtained through the Integrated Islet Distribution Program (IIDP) and the University of Alberta were enriched using a dithizone stain and cultured in CMRL 1066 supplemented with 10% FBS, 1X pen-strep, 8mM glucose, 2mM L-glutamine, 1mM sodium pyruvate, 10mM HEPES, and 250ng/mL Amphotericin B. For cytokine-treated samples, human cytokines were added to the culture media for 24 hours as follows: for high doses, 10ng/mL and IFN-g, 0.5ng/mL IL-1B (2 cytokines), with 1ng/mL TNF-α where indicated (3 cytokines); for low doses, 0.2ng/mL IFN-g and 0.01ng/mL IL-1B (2 cytokines), with 0.02ng/mL TNF-α where indicated (3 cytokines). Islet studies were approved by the Institutional Review Board of the University of California San Diego.

### Islet nuclei isolation

Human islets were collected from culture and homogenized in permeabilization buffer consisting of 5% BSA, 0.2% IGEPAL-CA630, 1mM DTT, and 1X cOmplete EDTA-free protease inhibitor (Sigma) in 1X PBS. Isolated nuclei were resuspended in 1X TDE1 buffer (Illumina) and quantified using a Countess II Automated Cell Counter (Thermo).

### ATAC-seq data generation

Approximately 50,000 nuclei were tagmented in a 25uL reaction volume using 2.5uL TDE1 (Illumina). Transposition reactions were carried out for 30 minutes at 37C in a thermal cycler. Tagmentation reactions were cleaned up using a 2X reaction volume of Ampure XP beads (Beckman Coulter) and used to prepare libraries using the Nextera XT Dual-Indexed primer system (Nextera) and NEBNext High-Fidelity PCR Master Mix (New England Biolabs). Libraries were sequenced by the UCSD Institute for Genomic Medicine on an Illumina HiSeq 4000 using paired end reads of 100bp.

### ATAC-seq data analysis

#### Processing

FASTQ reads were trimmed using Trim Galore (https://www.bioinformatics.babraham.ac.uk/projects/trim_galore/) with flags ‘--paired’ and ‘--quality 10’ and aligned to the hg19 reference genome with BWA mem^67^ using the ‘-M’ flag. We used Picard to mark duplicate reads and filtered, sorted, indexed, and aligned reads using samtools^68^ with flags ‘-q 30’, ‘-f 3’, ‘-F 3332’. Mitochondrial reads were also removed. Peaks were called on the filtered reads using MACS2^69^ with parameters ‘--extsize 200 --keep-dup all --shift - 100 --nomodel’. We generated bigWig tracks normalized by RPKM for each experiment using bamCoverage^70^. TSS enrichment scores for each ATAC-seq experiment were calculated using ‘tssenrich’ (https://pypi.org/project/tssenrich/), as the aggregate read distribution in a 4kb window centered on the TSS and normalized to an extended region of 1.9 kb on each side, according to the Encyclopedia of DNA Elements (ENCODE) guidelines.

#### PCA

We identified all peaks identified in at least two individual samples and constructed a read count matrix using edgeR^71^. We then calculated normalization factors using the ‘calcNormFactors’ function and used limma to apply the voom transformation and regress out batch effects and sample quality as measured by TSS enrichment scores. We then calculated principal components (PCs) using the top 10,000 most variable peaks using the ‘prcomp’ function with rank 2. The software used to generate PCs is located at https://rdrr.io/github/anthony-aylward/exploreatacseq^72^.

#### Differential chromatin accessibility

We generated a ‘master’ set of consensus ATAC-seq peaks by merging reads from all experiments and calling peaks on these merged reads using MACS2 as described above. The peaks were filtered to remove sites found in less than three individual samples and the ENCODE hg19 blacklist v2^73^. A count matrix of reads from each sample mapping to this list of peaks was created using featureCounts^74^ and used for differential accessibility analysis using DESeq2^12^. We used the experimental design ‘~treatment + donor’ and a cutoff of FDR<0.1 as computed by the Benjamini-Hochberg method to call differentially accessible sites between treated and untreated conditions. To compare the effects of treatment with and without TNF-α, we compared the absolute log_2_ fold changes from DESeq using a Wilcoxon signed rank test in R. To identify differentially accessible sites with different effects at different treatment durations we performed a linear regression of log2 fold changes with respect to matched controls as a function of time (6, 24, 48 and 72 hours). A nominal p-value of 0.01 was chosen as threshold.

#### Motif enrichment analysis

We used the ‘findMotifsGenome’ tool from HOMER^13^ to test differentially accessible chromatin sites for motif enrichment compared to a background of consensus ATAC-seq peaks, and using the masked hg19 genome as reference.

### RNA-seq data generation

RNA was isolated using the RNeasy Mini system from Qiagen from a total of 16 samples of human islets from 4 different donors and exposed for 24 hours to either 3 cytokines-high dose, 2 cytokines-high dose, 3 cytokines-low dose, 2 cytokines-low dose or control conditions. Approximately 500-1000 islets were used per sample. RNA quality was assessed using a 2200 TapeStation to confirm RNA integrity, and all samples had a RINe score of >7. Ribodepleted total RNA libraries were prepared and sequenced by the UCSD Institute for Genomic Medicine on an Illumina HiSeq 4000 using paired end reads of 100bp.

### RNA-seq data analysis

We used STAR (2.5.3a)^75^ to align paired-end RNA-Seq reads to hg19 genome with a splice junction database built from the Gencode v19 gene annotation^76^ and the following parameters: --outFilterMultimapNmax 20 --outFilterMismatchNmax 999 --alignIntronMin 20 --alignIntronMax 1000000 --alignMatesGapMax 1000000 --outSAMtype BAM Unsorted --quantMode TranscriptomeSAM. Gene expression values were quantified using the RSEM package (1.3.1) ^77^ with default parameters and loaded into R for further processing. Genes were filtered for >0.1 TPM on average per sample with 22,175 genes remaining after filtering. Raw expression counts were normalized using *voom* transformation from *limma* package and corrected for sample batch effects using limma removeBatchEffect. The R prcomp function was used to perform principal component analysis for the top 500 most variable genes. We identified differentially expressed genes between each cytokine treatment (3 cytokines-high dose, 2 cytokines-high dose, 3 cytokines-low dose, 2 cytokines-low dose) and untreated conditions using DESeq2^12^ with default settings and controlling for sample of origin using design= ~ sample + condition. An FDR of 10% was chosen as significance threshold. Metascape (https://metascape.org) was used to perform gene ontology enrichment analysis with standard settings.

### Single nuclei ATAC-Seq data generation

Islet nuclei from 4 donors (3 samples treated for 24h with 3 cytokines, high doses, and 4 untreated samples) were prepared as described above, and adjusted to a concentration of approximately 3000 nuclei/uL. Samples from two donors and same treatment conditions (SAMN12833535 and SAMN12889245) were pooled prior to snATAC library preparation and were de-multiplexed after sequencing as described below. We targeted 5000 nuclei per assay for use in the 10X Genomics Chromium Single Cell ATAC assay using v1 chemistry. Sequencing was performed at the UCSD Institute for Genomic Medicine on an Illumina NovaSeq using a specific paired-end 10X ATAC run configuration with reads of 50bp to a final read depth between 49 to 109M reads per sample.

### Single nuclei ATAC-Seq data analysis

#### Processing

10x Genomics Cell Ranger ATAC v1.1 (cellranger-atac count) was used to process 10x fastq files for each sample and perform alignment to the hg19 reference genome. For each assay, we then removed barcode multiplets using Cell Ranger’s multiplet removal script (version 1.1). BAM files were filtered for PCR duplicates, converted into tagAlign files, and intersected with a reference set of islet ATAC-seq peaks^78^ to construct a sparse matrix containing read counts in peaks for each cell. Cells with a minimum of 500 (sample SAMN15337453, untreated), 1,000 (samples SAMN12833535 and SAMN12889245), or 4,000 (sample SAMN15337453, treated and sample SAMN15314807) total mapped reads were retained for further analysis.

#### Clustering

Prior to combining all samples, each assay was clustered separately using scanpy v.1.6.0^79^. First, we extracted highly variable peaks using mean read depth and dispersion. Read depth was normalized and log-transformed counts were regressed out within highly variable peaks. We then performed PCA analysis and obtained the top 50 principal components. We calculated the nearest 30 neighbors using cosine metric to perform UMAP dimensionality reduction (min_dist = 0.3) and clustering using the Leiden algorithm. For each assay, cells with low usable counts and fraction of reads in peaks were iteratively removed. In order to obtain more accurate clustering and cell type assignment, the filtered assays were then merged and combined with 3 existing islet snATAC datasets previously filtered using the same criteria as above^80^, and the top 50 PCs were obtained from the merged experiments. Harmony^81^ was then used to batch correct PCs for donor across experiments. Using the corrected PCs, we applied the UMAP dimensionality reduction method, and clustered cells using the Leiden algorithm (Resolution = 0.5), and sub-clustered using the Louvain algorithm (Resolution = 1.5). Low-quality cells from the merged clusters were iteratively removed and manual doublet removal was performed on sub-clusters with above average high usable read depth or those that expressed multiple marker genes. After the entire filtering process, 28,853 cells were removed in total, and the final merged cluster contained 25,200 cells (untreated cells: 21,318; cytokine-treated cells: 3882) mapping to 10 clusters. Cell type of each cluster was assigned based on chromatin accessibility at promoter regions of known marker genes^78^ and verified through UCSC genome browser tracks. The islet samples from Wang et al^80^ were removed from the final clustering and the remaining cells (untreated cells: 3,947, cytokine-treated cells: 3,882) were used for downstream analysis.

#### snATAC pooled sample demultiplexing

In order to assign the pooled assays to the two sample donors, we genotyped non-islet tissue from the two samples. During islet picking, non-islet cells were collected separately from the islets, washed with 1X HBSS, pelleted at 500rcf for 5 minutes, and snap frozen with liquid nitrogen until genomic DNA extraction. We extracted genomic DNA from the non-islet cells using the PureLink Genomic DNA mini kit. Samples were genotyped by the UCSD Institute for Genomic Medicine using the Illumina Infinium Omni 2.5-8 assay. Genotypes were called using GenomeStudio (v2.0.4) with default settings. Using PLINK^82^, we filtered out rare variants with MAF <0.01 in the Haplotype Reference Consortium panel r1.1 and ambiguous alleles with MAF > 0.4. Filtered variants were used to impute genotypes into the HRC r1.1 panel using the Michigan Imputation Server with minimac4. Genotypes with high imputation quality (R2>0.3) were used to demultiplex pooled snATAC samples using Demuxlet^83^ with default settings.

#### Peak Calling

To identify chromatin accessibility peaks in each islet cell type, we extracted the reads from all cells within a given cluster and generated separate tagAlign files for each cell type. To correct for the 9-nt duplication created by Tn5 transposase, we shifted the reads aligned to the positive strand by +4bp and reads aligned to the negative strand by −5bp. We then called peaks using MACS2^69^ with the parameters ‘q 0.05’, ‘--nomodel’, ‘--keep-dup all’, and ‘g hs’. Blacklisted regions (v.2) from ENCODE were removed. The bedgraph output by MACS2 was sorted, normalized to counts per million (CPM), and converted to bigwig for visualization on UCSC genome browser. The peak calls from the individual cell types were then used to annotate the consensus set of peaks identified in bulk islet ATAC using bedtools intersect (v2.26.0).

#### Differential chromatin accessibility in islet cell types

We generated distinct BAM files for each cell type, donor and condition, using the barcodes to extract reads from the filtered and duplicate-removed BAM files from each assay using ‘samtools’ and ‘grep’. For each cell type, we then generated a matrix of read counts mapping to bulk ATAC consensus peaks using featureCounts^74^. Each matrix was filtered for an average read depth of 1 per sample/condition and DESeq2 was used to identify differentially accessible sites between cytokine treated and untreated samples with the donor as covariate (design = ~ treatment + donor). FDR <0.1 was chosen as significance threshold. To visualize results as heatmap and hierarchical clustering, we concatenated the matrices for each cell types, normalized the raw counts using DESeq *variance stabilizing transformation* (*vst*) function, filtered for peaks with differential accessibility in at least one cell type and plotted the resulting matrix using ‘pheatmap’. To compare the effects of cytokine treatment between beta cells and bulk islets and between beta cells and alpha cells, we compared the absolute log_2_ fold changes from DESeq at the same peaks using a Wilcoxon signed rank test in R.

#### Co-accessibility

Using Cicero (version 1.4.4)^15^, we calculated co-accessibility between pairs of snATAC peaks. To indicate which cells were accessible in which peak, we created a sparse *m* x *n* binary matrix by encoding cells from a given cell type (*n*) and merged peaks across all cell types (*m*), obtained using bedtools merge. We calculated Cicero co-accessibility scores following the recommended analysis protocol (https://cole-trapnell-lab.github.io/cicero-release/docs/#recommended-analysis-protocol), using the 30 nearest neighbors of UMAP coordinates to aggregate cells, and a window size of 1Mb to calculate cicero models. We then set a threshold of 0.05 and a minimum distance of 10kb to define pairs co-accessible for a given cell type. Co-accessibility was calculated for either untreated beta cells, cytokine-treated beta cells or merged treated-untreated beta cells. To annotate co-accessibility links between distal and promoter peaks, we categorized peaks within a 5kb window of a transcription start site (+/− 2.5 kb from TSS (GENCODE version 19^76^) as ‘promoter’, and otherwise as ‘distal’. To calculate enrichment in cytokine responsive cCRE for concordant effects with distal genes, we annotated each bulk ATAC consensus peak with results of differential accessibility in islets (3 cytokines, high-doses, 24 hrs) and co-accessibility in beta cells (merged treated-untreated conditions) with at least one gene with differential expression (3 cytokines, high-doses, 24 hrs). We then performed Fisher’s exact test on each combination of direction of effects (upregulated cCRE vs upregulated gene, upregulated cCRE vs downregulated gene, downregulated cCRE vs upregulated gene and downregulated cCRE vs downregulated gene). The same test was performed for cCREs proximal to gene promoters (<10 kb from TSS).

#### Motif enrichment analysis

Using ChromVAR^14^ (version 1.8.0) we calculated the deviation in accessibility from expected accessibility within islet cell types. We used a binary sparse matrix of accessible cells in each ATAC peak (see above) as input, and add GC bias using the ‘BSgenome.Hsapiens.UCSC.hg19’ library for genome sequence input. We then filtered cells with a minimum depth of 1500 and a minimum proportion of reads in peaks of .15, and filtered peaks for non-overlapping coordinates. The remaining peaks were annotated for motif occurrence from the JASPAR database, using the matchMotifs function from the motifmatchr package. We then computed deviations and variability for each cell type separately (alpha and beta) with the provided ChromVAR functions. For each transcription factor (n=386) we then calculated the absolute difference of the average deviation scores of cytokine treated and untreated cells and compared these differences in alpha versus beta cells using a scatterplot.

### EndoC-βH1 cell culture

EndoC-βH1 cells were cultured at 9×10^4^ cells/cm^2^ of cell culture surface area pre-coated with ECM (Sigma, E1270) and Fibronectin (Sigma, F1141). Cell culture media containing DMEM (Gibco,11885084), 2% BSA (Sigma, A1470), 3.5× 10^−4^% 2-mercaptoethanol (Gibco, 21985023), 0.12% Nicotinamide (Calbiochem, 481907), 5.5 ng/mL transferrin (Sigma, T8158), 6.7pg/mL Sodium Selenite (Sigma, 214485) and 1% Penicillin-Streptomycin (Gibco, 15140122) were refreshed every 2 days. Cells were passaged weekly using 0.25% Trypsin-EDTA for dissociation, which was quenched with an equal volume of FBS and two volumes of IMDM media (Gibco, 12440053). Dissociated cells were spun down at 1200 rpm for 5 minutes and counted before seeding with the above-mentioned density.

### HiChIP sample preparation

To collect samples for HiChIP assays, 10 million EndoC-βH1 cells were treated with either control (0.1% BSA) or cytokines (0.5 ng/mL IL1β, 1 ng/mL TNFα and 10 ng/mL IFNγ) for 72 hours. Treated cells were cross-linked with 1% formaldehyde for 15 minutes with shaking at room temperature, followed by a 5-minute quenching step with 1.25 M glycine/PBS. Cross-linked EndoC-βH1 cells in both control and cytokine-treated conditions were washed three times with ice-cold PBS and collected from the dish with a cell scraper. Cells were then pelleted, and flash frozen with liquid nitrogen. HiChIP assays were performed by Arima Genomics using the HiC+ protocol with a H3K27ac antibody, and libraries were sequenced on an Illumina NovoSeq with 150 bp paired end reads.

### HiChIP data analysis

Data was processed using the MAPS v2.0 pipeline from Arima Genomics (https://github.com/ijuric/MAPS/tree/master/Arima_Genomics). We used hg19 as the reference genome and H3K27ac ChIP-seq peaks in EndoC-βH1 cells from a published study^84^. Interactions between 5kb windows containing a H3K27ac peak were considered significant at FDR<0.10. Loop calls were intersected with promoter regions of genes from GENCODE to identify enhancer-promoter interactions. Contact matrices were generated using the *pre* command from Juicer tools^85^. For virtual 4C, we extracted all contacts which included the 5kb window around the site of interest.

### Lentiviral human GeCKO-V2 library preparation, transduction, and titration

To package lentivirus encoding the human GeCKO-V2 CRISPR screen library^86^, plasmids containg the gRNA library (Addgene, 1000000048) were transfected into the HEK 293T cells together with the lentiviral packing vectors, pMD2.G (Addgene, 12259) and psPAX2 (Addgene, 12260), using a PolyJet™ DNA transfection reagent (Signagen Laboratories, 504788). Transfected cells were kept in the culture to allow virus to be released. And the media containing lentivirus was collected at 36, 48, 72 hours post transfection before filtered through a 0.45μm cell strainer to remove cell debris. Lentiviral particles were pelleted down at 20,000 rcf for 2 hours, using an Optima L-80 XP Ultracentrifuge machine (Beckman Coulter) provided by the Human Stem Cell Core at UCSD. The same media for EndoC-βH1 cell culture was used to resuspend the virus.

A spin-inoculation method was adopted to transduce the viral library into the EndoC-βH1 cell line. To do this, the cells were pre-treated with 8 μg/mL polybrene (Sigma, TR-1003) in the culture media for 30 minutes. Then the virus was added before the entire plate was spun in a swing-bucket centrifuge machine at 930g for 45 minutes. It takes 48 hours for the sgRNA and Cas9 protein to be expressed in the EndoC-βH1 cells.

### CRISPR loss-of-function screen for regulators of β-cell survival under cytokine stress

The EndoC-βH1 cells were expanded to a total of 300 million cells before spin-inoculated with the lentiviral human GeCKO-V2 library at an MOI=0.3. To enrich for successfully transduced cells, a 3-day puromycin (5 μg/mL, Sigma, P8833) selection was performed 48 hours after the spin-inoculation. And 60M (500X genome coverage) cells were harvested as a representation control for the GeCKO-V2 sgRNA library. The rest of the cells were kept in the culture condition for an additional 14 days with puromycin (1 μg/mL) to achieve sufficient gene deletion and treated with either 0.1% BSA or a combination 0.5 ng/mL IL1β (PerroTech, 200-01B), 1 ng/mL TNFα (PerroTech, 300-01A) and 10 ng/mL IFNγ (PerroTech, 300-02) for 72 hours. A time-point experiment was performed to evaluate which treatment duration was necessary to induce cell death. EndoC-βH1 cells were seeded 24 hours before the cytokine treatment and residual cell number was counted at 24, 48 and 72 hours of treatment (n=3). Cell number are shown in **Supplementary Table 5**. We were able to harvest another 60M (500X genome coverage) cells from the control (0.1% BSA) treated cells and 30M (250X genome coverage) from the cytokine treated cell, although they were started with the same number.

Genomic DNA from all three conditions were purified with a Quick-gDNA™ MidiPrep kit (Zymo Research, D3100). And sgRNA library were amplified from the genomic DNA using a two-step nested PCR method modified from a previous published protocol (PMID: 28417999). In brief, guide RNA inserts were amplified from the genomic DNA with the following primers:

F1-1:TCCCTACACGACGCTCTTCCGATCTNNNNNGGAAAGGACGAAACACCG
F1-2:TCCCTACACGACGCTCTTCCGATCTNNNNNHGGAAAGGACGAAACACCG
F1-3:TCCCTACACGACGCTCTTCCGATCTNNNNNHHGGAAAGGACGAAACACCG
F1-4:TCCCTACACGACGCTCTTCCGATCTNNNNNHHYGGAAAGGACGAAACACCG
R1-1:GGAGTTCAGACGTGTGCTCTTCCGATCNNNNNTGCTATTTCTAGCTCTAAAAC
R1-2:GGAGTTCAGACGTGTGCTCTTCCGATCNNNNNVTGCTATTTCTAGCTCTAAAAC
R1-3:GGAGTTCAGACGTGTGCTCTTCCGATCNNNNNVMTGCTATTTCTAGCTCTAAAAC
R1-4:GGAGTTCAGACGTGTGCTCTTCCGATCNNNNNVMAATGCTATTTCTAGCTCTAAAAC

Pooled F1 primers (F1-1 to F1-4) and R1 primers (R1-1 to R1-4) were used each PCR reaction to avoid cluster registration failure on Illumina machines. Amplicons from the first step of PCR were gel purified and subjected to a second round of PCR to add Illumina sequencing adaptors and TruSeq indexes. Primers used in the second PCR step were listed below:

F2:AATGATACGGCGACCACCGAGATCTACACTCTTTCCCTACACGACGCTCTTCCGA
R2:CAAGCAGAAGACGGCATACGAGATNNNNNNGTGACTGGAGTTCAGACGTGTGCTCTTC CG

Sequencing library amplified from the second round of PCR were size-selected and purified with a magnetic bead-based SPRIselect reagent (Beckman Coulter, B23318), and subjected to HiSeq4000 Illumina NGS platform using a single read (SR75) method.

### Analysis of CRISPR screen results

Adaptor sequences ggaaaggacgaaacaccg and gttttagagctagaaatagca flanking the 19-20 base pair of sgRNA sequences were trimmed using cutadapt. Trimmed sequencing reads were then aligned to the reference sgRNA library with bowtie2 with default settings, resulting in a BAM file that can be used for sgRNA counting with the MAGeCK model-based tool for CRISPR-Cas9 knockout screens^87^. Statistical significance of guide RNA representation in control and cytokine treated datasets was estimated with the mle subcommand in the MAGeCK package. An effect (beta) for each gene from this analysis was extracted as an indicator of enrichment (positive beta) or depletion (negative beta) of sgRNAs targeting this gene in the cytokine-treated cells. miRNA genes and genes with less that 3 sgRNA guides were excluded from further analysis. Significantly enriched (427) or depleted (440) genes at FDR<.10 were further filtered for expression in islet (average sample TPM>=1). Gene ontology analysis was performed using GSEA (http://www.gsea-msigdb.org/gsea/msigdb/annotate.jsp) against the REACTOME and GO biological process gene sets, including only gene sets with more than 20 and less than 1000 genes. A Fisher’s exact test was used to calculate the enrichment of pro-cell survival and pro-cell death genes segregated by up-regulated, down-regulated or no change in expression in high-dose cytokine treatment within 1Mb of all known T1D risk loci including MHC.

### SNP-SELEX variant selection

Variants were selected and classified based on 4 criteria. (1) *T1D loci:* We selected 86,067 variants from 57 known T1D loci, including the MHC region. Variants at these 57 loci were selected based on: credible set variants from fine mapping data for 36 loci^88^, all variants in 1,000 Genomes Project (1KGP) phase 3 EUR LD (r2>0.2) with index variants at the remaining 21 loci, and all variants in 1KGP with EUR MAF >0.5% in regulatory elements within 250 kb of index variants at all 57 loci. (2) *T2D* loci: We selected 33,354 variants at known T2D loci, which include lead variants and variants in LD with r2>=0.6 in EUR and non-EUR, and credible variants from fine mapping studies. (3) *Islet enhancers:* We included 56,796 variants in 1KGP phase 3, filtered for Hardy-Weinberg Equilibrium p-value >=1e-5 and MAF >=0.5% that intersected with islet enhancers, defined using published ATAC-Seq and H3K27ac ChIP-Seq data from human islets^89,90^. (4) *Random*: 7,869 negative control variants from filtered 1KGP SNPs, but randomly chosen from the genome were included. Variants from categories 2, 3 and 4 have been included as a validation set in a previous publication^26^. The total number of selected variants is 184,086, including 183,373 SNPs and 713 indels. A small subset of variants overlaps between the 4 different selection methods, and therefore in total there were 182,226 distinct variants selected.

### SNP-SELEX data generation

Oligonucleotide design consisted of a target sequence of 44 nt containing the variant, flanked by illumina TruSeq dual-index system adapters and barcodes. Three hundred and eighty-four pools of oligonucleotides were synthesized by CustomArray (Seattle, WA), each pool carrying a unique sequence barcode. To control for PCR duplicates, the 3 nucleotides at each end of the 44 nt sequence were synthesized Ns, which generated random combination of nucleotides tagging each molecule. For SNPs, the central position was substituted by an N, resulting in synthesis of all 4 nucleotidic variations (97,758 oligos), while for indels (maximum 3bp-long) both a long (44 nt) and a short (41-43 nt) form were synthesized (259 x 2 oligos). The oligos were double stranded using 20 cycles of PCR and sequenced for 2×50 paired-end cycles with illumina Hiseq 2500 as input references.

SNP-SELEX data was generated as previously described^26^. The cDNA of 530 distinct TF proteins were cloned into pET20a plasmids and expressed using Rosetta (DE3) pLysS *E. coli* strains. 6xHis-tagged TF proteins were immobilized to Ni sepharose beads (GE, 17-5318-01) in Promega binding buffer (10mM Tris pH7.5, 50mM NaCl, 1mM MgCl2, 4% glycerol, 0.5mM EDTA, 5μg/ml poly-dIdC) across 8×96-well plates. Oligos from input were added into the protein beads mixture and incubated at RT for 30 min. Beads were washed for 12 times with the Promega binding buffer and re-suspended in TE (10mM Tris pH 8.0, 1mM EDTA). The eluted DNA was amplified by PCR and purified (Qiagen, 28004): an aliquot used for library preparation and sequencing and another aliquot of the same product was added to the protein beads mixtures for a new binding cycle. Two independent replicates consisting of four binding cycles each were performed and sequenced using two flow cells of 2×50 paired end illumina Hiseq 2500. To reduce confounders due to systematic synthesis bias, in the second experiment the order of the input pools was inverted (i.e. the same TF protein was hybridized to an oligo pool synthesized with a different barcode).

### SNP-SELEX sequencing data analysis

#### Sequence processing

FASTQ files from each cycle and input were first filtered for identical sequences using FastUniq (v1.1)^91^, which removed on average 10% of reads in each experiment, to a final median depth of 3 and 0.64 million paired-end reads for the input and the selected oligos respectively. Sequencing reads were then aligned using BWA-MEM (version 0.7.12)^67^ to the oligo library fasta files. For each oligo, the number of read pairs carrying each nucleotide was counted, only counting reads that were uniquely mapped, correctly paired, with quality=60 and with the same sequence at the SNP position. Oligos with less than 8 read pairs for SNPs and 4 reads pairs for indels were excluded for further analysis. To estimate the consistency between the two experimental replicates we calculated the correlation between the proportion of reads aligning to a given SNP over the total number of reads in a given experiments. The proportion of reads aligning to each oligo in each experiment was well correlated between the two replicates (median Pearson coefficient r=0.86), and the correlation increased over the cycles, indicating that the selection for the same oligo by a given protein was reproducible between the two replicates. We excluded that this result was due to the starting oligonucleotide stoichiometry in the pools, as the different replicates had different input material.

Motif analysis was performed on the oligo sequences (40 nt) that were selected at cycle 4 of each experiment to determine the enrichment for the expected motif or family of motifs. For motif enrichment we used HOMER (library 4.7)^13^ and MEME 4.12.0 (libraries: JASPAR_CORE_2014_vertebrates, jolma2013, encode_known, Mariani_2017 and Barrera_2016)^92^. A positive motif match was determined if the expected motif (matching with the first three letters of the name) or a motif from the same structural family (defined by homer classification) were found among the top 20 enriched motifs. For 564 experiments, we found a positive motif match in both replicates, for 90 in either of the two replicates and for 114 in none of the replicates. Because for some analyzed TFs the motif is not known, for example for Zinc Finger proteins, we did not consider failed experiment only based on the motif enrichment, but also on the correlation between replicates. If the correlation between replicates was <0.5 and one of the two replicates was enriched for the expected motif, then remove only the replicate that did not contain the motif (16 experiments removed). If the correlation was <0.5 and both replicates did not have motif in the corresponding family, we removed both replicates (53 experiments removed).

#### Identification of variants with differential TF binding

For each experimental replicate, allelic counts were tabulated for each oligo at each cycle, including only those variants covered by at least 8 read pairs for SNPs, or 4 reads pairs for indels, in all five cycles (0-4). Furthermore, variants with less than 2 read pairs in the input for both the reference and alternate alleles and composing < 5% of the total reads in the pool were removed, as potentially biased inputs. To quantifying the magnitude of the difference between reference and alternate allele binding across all cycles, we used the “Preferential Binding Score” or PBS, which has been previously described^26^. The PBS corresponds to the AUC between the differences of log odds ratios of the two alleles compared to cycle 0 (the input), and is calculated as follows:

1. For a given oligo, the odds of allele *a* at cycle *c* is defined by the frequency *P* of allele *a* at cycle *c*, divided by 1-*Pa,c*, which is equal to the read counts of allele *a* divided by the sum of read counts of all other nucleotides *r*: *Odds a,c = P a,c / (1-P a,c) = counts (a,c) / counts(r,c)*
2. The odds ratio is calculated as the ratio between the odds of allele *a* at cycle *c* and the odds of allele a at cycle 0: *OR a,c = Odds a,c / Odds a,0*
3. LogOR are calculated for reference and alternate allele for each cycle: *LogORa,c = log10(counts (a,c)) + log10(counts (rest, 0)) - log10(counts(a,0)) −log10(counts (rest,c))*
4. The PBS is the AUC of the difference between LogORref and LogORalt (∆LogOR), calculated with the formula: 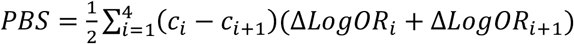

For each experiment replicate, to determine the statistical significance of the observed values, a Monte Carlo randomization was conducted, which consisted of 250,000 randomly generated PBS measurements. The randomizations consisted of shuffling the SNP labels 250,000 times within each cycle and one PBS measurement was extracted each time. We observed that experiments with fewer than 25 oligos generated non-normal PBS random distributions, therefore experiments with less than 25 variants remaining after the above filtering steps were excluded.

After calculating preferential binding statistics in each individual experiment (same “well”, two technical replicates), results of the two replicates for each experiment were combined using meta-analysis of p-values, weighted on the total number or reads for reference and alternate allele in cycles 1 to 4, and the average of effect sizes (PBS). Further, experimental replicates of the same TF protein (different “wells”, variable number of replicates) were meta-analyzed to obtain a unique value for each TF. A nominal p-value of 0.05 was chosen as arbitrary threshold to define a preferentially bound variant.

#### Correlation of SNP-SELEX results between transcription factors

To compare variant effects on binding of different TFs, we first computed a matrix of PBS scores where each row corresponded to a SNP and each column to a TF. After filtering the matrix to retain only TFs with at least 50 bound variants, TF families with at least 3 components and variants that were pbSNPs in at least one TF (27,655 variants and 457 TFs), we calculated a pairwise correlation matrix using the cor() function in R, using the “pairwise” option. To perform hierarchical clustering on TFs, we filtered the pairwise correlation matrix, retaining only rows and columns with non-missing values (264 TFs). The dendrogram of the hierarchical clustering of distances was obtained using the R command as.dendrogram(hclust(as.dist(1-correlation_matrix))) and plotted using functions form the “circlize” R package.

#### Correlation between SNP-SELEX and SNP effect predictions

Predictions of TF motif alteration by SNPs were calculated using the package motifbreakR^93^, using the *H.sapiens* ‘HOCOMOCOv10’ library of motifs PWM (640 TFs, including most of those tested by SNP-SELEX) and using the parameters: filterp = TRUE, method=“ic”, threshold = 5e-4, BPPARAM = BiocParallel::bpparam(“SerialParam”). We tested using this approach 181,540 variants (some where excluded in the formatting process by the function: snps.from.file (search.genome = BSgenome.Hsapiens.UCSC.hg19), of which 177,270 were predicted to alter at least one of the motif of the library. For each SNP, the difference of PWM scores from the two alleles was compared with the PBS scores from SELEX from the corresponding TFs (500 unique TFs, 129,842 unique variants, 1,896,977 combinations). Pearson correlation coefficient was calculated for 234s TF that had a minimum of 10 testable, bound SNPs with PWM predicted effects, or 146 when only considering pbSNPs. Similarly, predictions of SNP effect on TF binding were obtained from DeepSEA calculations (http://deepsea.princeton.edu/job/analysis/create/) and filtered for E-value <0.01. For each TF in the database, the predicted allelic log2 fold change of each SNP was averaged across the different cell types and then compared with SELEX PBS scores, for TFs having a minimum of 10 bound SNP (37 TFs: ATF2, ATF3, BATF, CEBPB, CTCF, E2F4, ELF1, ELK1, ELK4, ETS1, FOSL1, FOXA1, FOXA2, FOXM1, FOXP2, GATA2, GATA3, IRF3, IRF4, MEF2C, MYBL2, NANOG, NFATC1, NFIC, POU2F2, POU5F1, PRDM1, RFX5, RUNX3, RXRA, SRF, TCF12, TCF7L2, USF1, USF2, YY1, ZBTB7A) or pbSNPs (24 TFs).

#### Genetic association enrichment analysis

We tested variants with allelic effects on TF binding for enrichment of T1D association using genome-wide summary statistic data^8^. We defined three categories of variants: (i) all variants, (ii) mapping in beta cell cCREs, (iii) mapping in cytokine-responsive beta cell cCREs. For each variant category, we identified several different p-value thresholds and segregated SNP-SELEX variants based on (i) allelic effects of TF binding, or no allelic effect on TF binding, (ii) reaching p-value threshold or not, and then for each threshold performed a Fisher’s exact test.

### Electrophoretic Mobility Shift Assay

Electrophoretic Mobility Shift Assay (EMSA) was carried out using LightShift™ Chemiluminescent EMSA Kit (20148, ThermoFisher Scientific). Untreated and cytokine treated MIN6 nuclear extracts (NEs) were prepared using NE-PER Nuclear and Cytoplasmic Extraction Reagents as per manufacturer’s recommendation (78833, ThermoFisher Scientific), supplemented with 1x protease inhibitors (40694200, Roche Diagnostics GmbH). For cytokine-treated condition, MIN6 cells were cultured in T75 flasks to 70% confluency and treated with 10ng/mL IFN-γ, 0.5ng/mL IL-1β, and 1ng/mL TNF-α cytokine mixture prepared fresh 24 hours prior to NE preparation. NE protein concentration was determined using a NanoDrop (ThermoFisher Scientific) and samples were stored at −80°C until analyses. Sense and anti-sense single-stranded EMSA oligonucleotides for reference and alternate alleles were purchased from Integrated DNA Technologies, with the following sequences:

rs10483809 (*RAD51B*): 5’Biotin–ATCTTTCACTTTCCCT[***A/G***]TCGATACTTCATATGT
rs35342456 (*SOCS1*): 5’Biotin–GCTGGGCGTGGTGGCTCACGCCTGT**[A/C]**ATCTTGTTG

Binding reaction mixtures were prepared for each allele and contained 10x Binding Buffer, 50% glycerol, 0.1M MgCl_2_, 1μg/μL in 10mM Tris Poly(dI*dC), 1% NP-40 (20148, ThermoFisher Scientific), 100fmol and 25fmol of labeled probe for rs10483809 and rs35342456 respectively, and 8-17 μg NE. For corresponding competition reaction(s), 200-fold excess of unlabeled probe at (20 or 5 pmol) was used. Competition reactions were incubated at RT for 10 minutes with NE and unlabeled probe prior to adding biotin-labeled probe. Reaction mixtures were further incubated for 20 minutes at RT, and 5x Loading Buffer was added to each mixture to stop the reaction. Empty 6% TBE gel (EC62655BOX, Invitrogen) was run at 100V in 0.5x UltraPure TBE Buffer (15581-044, Invitrogen, Life Technologies) at 4 °C prior to loading samples. Samples were subsequently run on the same gel at 100V for 90 minutes at 4°C. DNA-protein complexes on the gel were transferred to 0.45mm Biodyne™ Pre-Cut Modified Nylon Membrane (77016, Thermo Scientific) at 380 mÅ for 45 minutes, and were crosslinked for 15 minutes using UV Transilluminator (VWR, VWR International). Complexes were detected using Chemiluminescent Nucleic Acid Detection Module (20148, ThermoFisher Scientific) after blocking for 1 hour. Images were captured using a C-DiGit Blot Scanner (Model 3600, Li-Cor Biosciences).

### Gene reporter assays

We cloned a 400bp insert containing the rs10483809 variant using human DNA from Coriell as a template into the pGL4.23 reporter vector in the forward direction using the restriction enzymes KpnI and SacI. A reporter containing the alternate allele was generated through SDM using the Q5 Site-Directed Mutagenesis kit (New England Biolabs). The primer sequences used were as follows:

rs10483809_cloning_FWD CCATGGTTTCTTCCTGGGTA
rs10483809_cloning_REV GCACAAAATAGAAGAAAGATCAAGAA
rs10483809_SDM_P1 TTTCTCTTTCgCAAACTCCTC
rs10483809_SDM_P2 TGTCACTGACTGAGTTGC

For gene reporter assays, MIN6 between passages 17-21 were plated at a density of 0.25E6 viable cells/mL in a 48-well plate. 500ng experimental vector was co-transfected with pRL-SV40 using Lipofectamine 3000 (Thermo Fisher). 48 hours post-transfection, cells were lysed and used in the Dual-Luciferase Reporter System assay (Promega). Firefly activity from the experimental plasmids, normalized by dividing by the corresponding Renilla activity, was compared to the normalized activity of the empty pGL4.23 vector. A two-sided t-test was used to compare the luciferase activity between the alternate and reference allele.

### DNA fragmentation ELISA

MIN6 cells between passages 17-21 were grown to approximately 80% confluency in 24-well plates and transfected with 30μM of either *Socs1* siRNA (Invitrogen Silencer select) or scramble siRNA (Invitrogen Silencer Negative Control No. 1) using the Lipofectamine 3000 agent (Invitrogen). 24 hours after transfection, cells were washed once with 1X PBS and labelled with 10μM BrdU for two hours. Cells were washed twice with 1X PBS before being given complete MIN6 media, with cytokines added to the indicated samples. 150uL of supernatant was collected at indicated times and spun down at 500rcf for 5 minutes to pellet cell debris. 100uL of the clarified supernatant was used in the anti-BrdU DNA fragmentation ELISA (Roche). Each condition was tested in technical triplicates, from MIN6 wells seeded from the same passage.

## ACKNOWLEDGEMENTS

The work performed in this study was supported by NIH grants DK122607 and DK120429 to KJG and MS, DK112155 to MS, KF, KJG and BR, and DK105541 to MS, KR and BR.

## AUTHOR CONTRIBUTIONS

KJG and MS conceived of and supervised the study. KJG, PB, HZ, MO, and MS wrote the manuscript. JT and BR supervised the design and generation of SNP-SELEX data. KF supervised the design of SNP-SELEX and the analyses of genomic data in the study. PB performed statistical data analysis of genomic, CRISPR-KO screen, and genetic data; HZ performed the CRISPR-KO screen and interpreted the results; MO performed bulk and single cell genomic assays. JY performed the SNP-SELEX assay. PB, MO, RE, EB, KK, AA, JC and JN performed bulk and single cell data analysis. MO, RE, JK and SC performed functional validation experiments. PB, NN, YQ, AA and MKD performed design and analysis of SNP-SELEX data. All authors contributed to and approved of the final version of the manuscript.

## DATA AVAILABILITY

The ATAC-seq, RNA-seq, snATAC-seq, HiChIP and CRISPR screen data will be deposited in GEO and CMDGA. The SELEX-seq data are in GEO under accession number GSE118725.

## Supplementary Material

**Supplementary Figure 1. Effect of different cytokine treatments on islet accessible chromatin.** A) Top: Scatterplot showing effect on cytokine-responsive cCREs (DESeq FDR <0.10) chromatin accessibility in islets after treatment with high doses of 3 cytokines (IL-1β, IFNγ, and TNFα , x-axis) versus 2 cytokines (IL-1β and IFNγ, y-axis). Bottom: density plot showing increased effect size in cytokine treatment with TNFα. Wilcoxon signed rank test p-value is shown. B) Heatmap of cytokine effect sizes (log_2_ fold change, DESeq) on cytokine-responsive cCREs that change with treatment duration (linear regression p< 0.01) C) Example of two cytokine-responsive cCREs at the *HEATR2* and *CRHR1* loci that show increased accessibility over time. D) Motif enrichment for up-regulated or down-regulated cytokine–responsive cCREs identified using different duration of cytokine treatments. Motifs that were significantly enriched in at least one condition (HOMER FDR<0.05, indicated by an asterisk) are shown. Red boxes highlight motifs with visible differences in enrichment over time.

**Supplementary Figure 2. Defining islet cell sub-types from snATAC-seq profiles.** A) UMAP plots showing clusters of islet snATAC. B) Proportion of cells derived from different donors in each cluster. C) UMAP plots showing promoter accessibility in a 1 kb window around the TSS for selected cell type marker genes. D) Genome browser plots showing aggregate read density (CPM-normalized read depth, range: 0-7, shown on vertical axis for each plot) for cells within each cell type for selected cell type marker genes.

**Supplementary Figure 3. Cell type-specific changes in islet accessible chromatin upon inflammatory cytokine exposure.** A) Number of cytokine-responsive cCREs (or DACs) in bulk islet ATAC that overlap a snATAC from different cell types. B) Number of DACs in bulk islet ATAC that overlap a snATAC specific to a cell type. C) Heatmap of z-score normalized chromatin accessibility at significant DACs (DESeq FDR<0.1) identified in beta, alpha and delta cells by snATAC comparing cytokine-treated and untreated samples. Endothelial, acinar and stellate cells did not show any significant DAC. C) Scatterplot showing DACs effect sizes (DESeq log_2_ fold change) in alpha and beta cells. D) Density plot showing increased cytokine response in beta cells at DACs significant in either beta or alpha cells (top), and in DACs significant in both cell types (bottom). E) Comparison of motif enrichment in chromatin accessibility from cytokine treated alpha and beta cells. ChromVAR deviation scores within alpha or beta cells were averaged across treated and untreated cells and their difference (∆CTY-UNT) was plotted in a scatterplot. The slope (ß) from linear regression and the most different motifs between alpha and beta are shown are shown.

**Supplementary Figure 4. Cytokine-induced gene expression changes in pancreatic islets.** A) Principal components plot of normalized and batch-corrected gene expression from high-dose-2-cytokine (orange), high-dose-3-cytokine (red) low-dose-2-cytokine (blue), low-dose-3-cytokine (green)-treated and untreated (purple) islets from a total of 16 samples. Donor ID is indicated on the top of each dot. B) Number of differentially expressed genes (DE genes, DESeq FDR<0.1) between each cytokine treatment condition and untreated islets. C) Venn diagram showing overlap between DE genes in each treatment. D) Heatmap showing the top 20 upregulated and top 20 downregulated genes common to each treatment vs untreated islets, and the top 10 differential genes (in bold) between high-dose-2-cytokine and high-dose-3-cytokine (i.e due to TNFα). E) Gene ontology terms enriched among genes with up-regulated expression in cytokine-treated islets. F) Gene ontology terms enriched among genes with down-regulated expression in cytokine-treated islets. G) Enrichment of islet distal differentially accessible cCREs (DACs) (>10kb from TSS) for genes with concordant cytokine-induced effects, linked by HiChIP (FDR <0.1). Fisher’s exact test p-values and odds ratios are shown. HiChIP was performed in untreated (left) or cytokine-treated (right) EndoC-βH1 cells.

**Supplementary Figure 5. SNP-SELEX sequencing metrics, replicate consistency, and comparison with TF binding predictions**. A) Fraction of reads retained after removing identical sequencing duplicate reads. The input is composed of 384 pools of oligos with different barcodes (from 4x 96-well plates); each SELEX cycle is composed of 768 assays (8x 96-well plates), performed twice. Median value is indicated at the top of each boxplot. B) Number of reads retained after removing identical sequencing duplicate reads. The y-axis is log scaled. C) Left: example of one experiment showing correlation between the percentages of reads mapping to each oligo (i.e. each dot) in replicate 1 versus replicate 2. Pearson correlation coefficient is indicated. Right: distributions of Pearson coefficients calculated as in the example, across all 768 experiments and cycles. D) Number of experiments showing enrichment at cycle 4 for motifs similar to the assayed TF protein in both replicates, only one of the two, or none. E) Hierarchical clustering of the pairwise distance (1-correlation) of allelic effects (PBS score) across different TF proteins, color-coded according to the structural family. 264 TFs that had a minimum of 100 testable SNPs are shown. F) Left: distribution of correlation between PBS and DeepSea Log fold change across TFs. The number of TFs analyzed (having both DeepSea predictions and SNP-SELEX results for at least 10 SNPs) are indicated. Rigth: scatterplot of PBS and DeepSea Log fold change across tested SNP-TF pairs (number indicated). pbSNPs are shown in purple. G) Top: distribution of correlation between PBS and ∆PWM across TFs. The number of TFs analyzed (having measurements for both PWMs and SNP-SELEX for at least 10 SNPs) are indicated. Bottom: scatterplot of PBS and ∆PWM across all tested SNP-TF pairs (number indicated). pbSNPs are shown in purple. H) Pearson correlation coefficients between SNP-SELEX PBS score and ∆PWM in each TF across all bound SNPs, grouped by structural families. 234 TFs that had a minimum of 10 testable SNPs with PWM predicted effects are shown. I) Percentage of pbSNPs that corresponded to a predicted PWM change in each TF, grouped by TF family. 234 TFs that had a minimum of 10 testable SNPs with PWM predicted effects are shown

**Supplementary Figure 6. Electrophoretic mobility shift assay (EMSA) for rs35342456 at the *DEXI/SOCS1* locus.** Three independent EMSA experiments (different cell cultures) and one replicate of binding reaction for experiment #3 are shown. MIN6 were cultured in control and cytokine media and nuclear extracts were used in binding reaction with oligonucleotides carrying either the reference (A) or alternate (C) allele of rs35342456. Both treated and untreated MIN6 cells nuclear extracts showed preferential binding to probes with the reference allele. The top-left panel (Experiment 1) shows the non-cropped image shown in Figure 6E.

**Supplementary Table 1.** Pancreatic islet donor samples

**Supplementary Table 2.** Cytokine-responsive cCREs in pancreatic islets and cell types

**Supplementary Table 3.** Sequence motifs enriched in cytokine-responsive islet cCRE and motif deviation between cytokine treated and untreated cells

**Supplementary Table 4.** Genes with differential expression in cytokine-treated islets

**Supplementary Table 5.** Genes affecting survival in cytokine-treated beta cells

**Supplementary Table 6.** GSEA for genes affecting survival in cytokine-treated beta cells

**Supplementary Table 7.** TFs tested in SNP-SELEX assay and motif enrichment

**Supplementary Table 8.** T1D risk variants in cytokine-responsive beta cell cCREs

