## Supplementary Figures for "Type 1 diabetes risk genes mediate pancreatic beta cell survival in response to proinflammatory cytokines"

**Supplementary Figure 1**

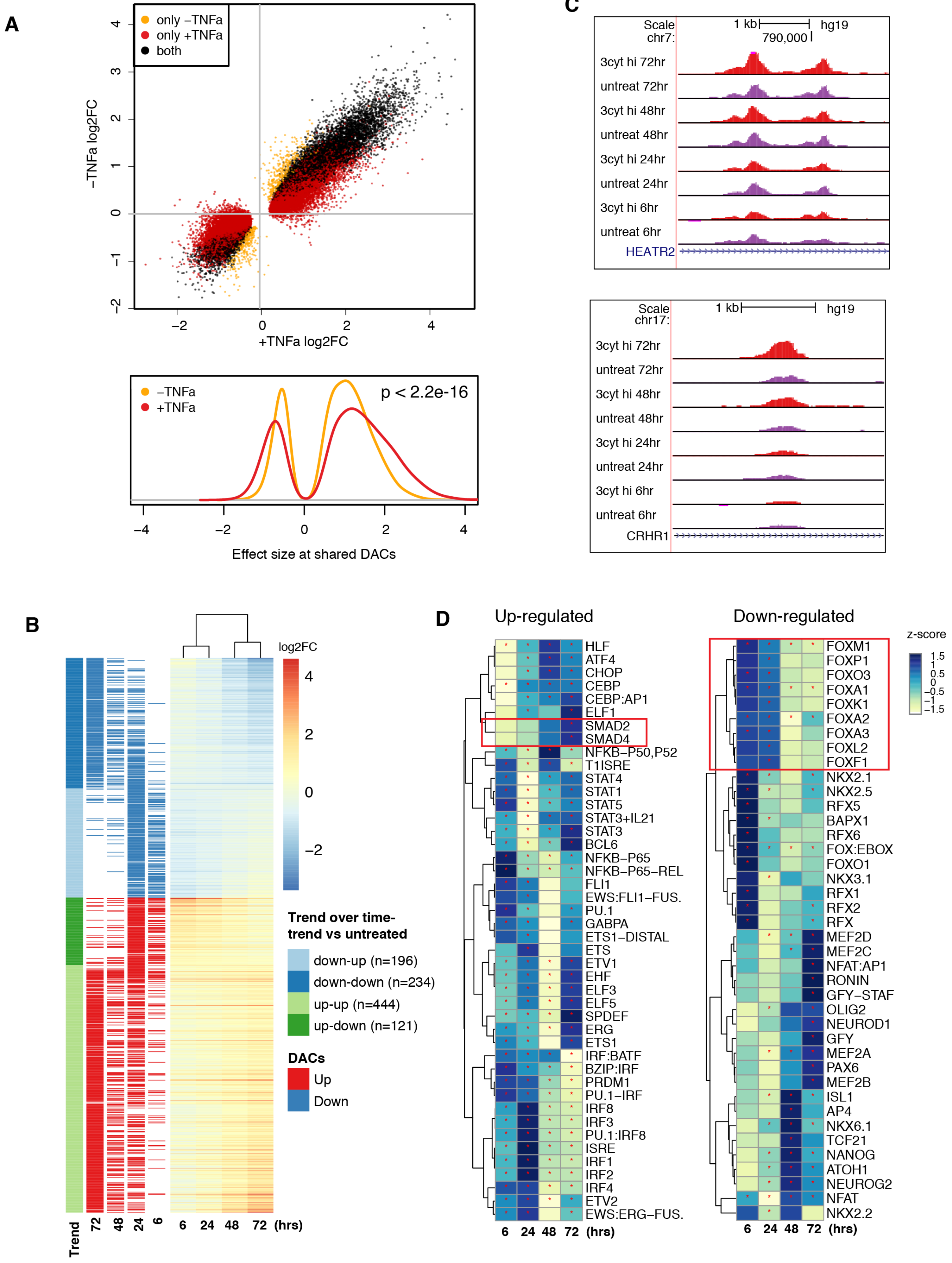

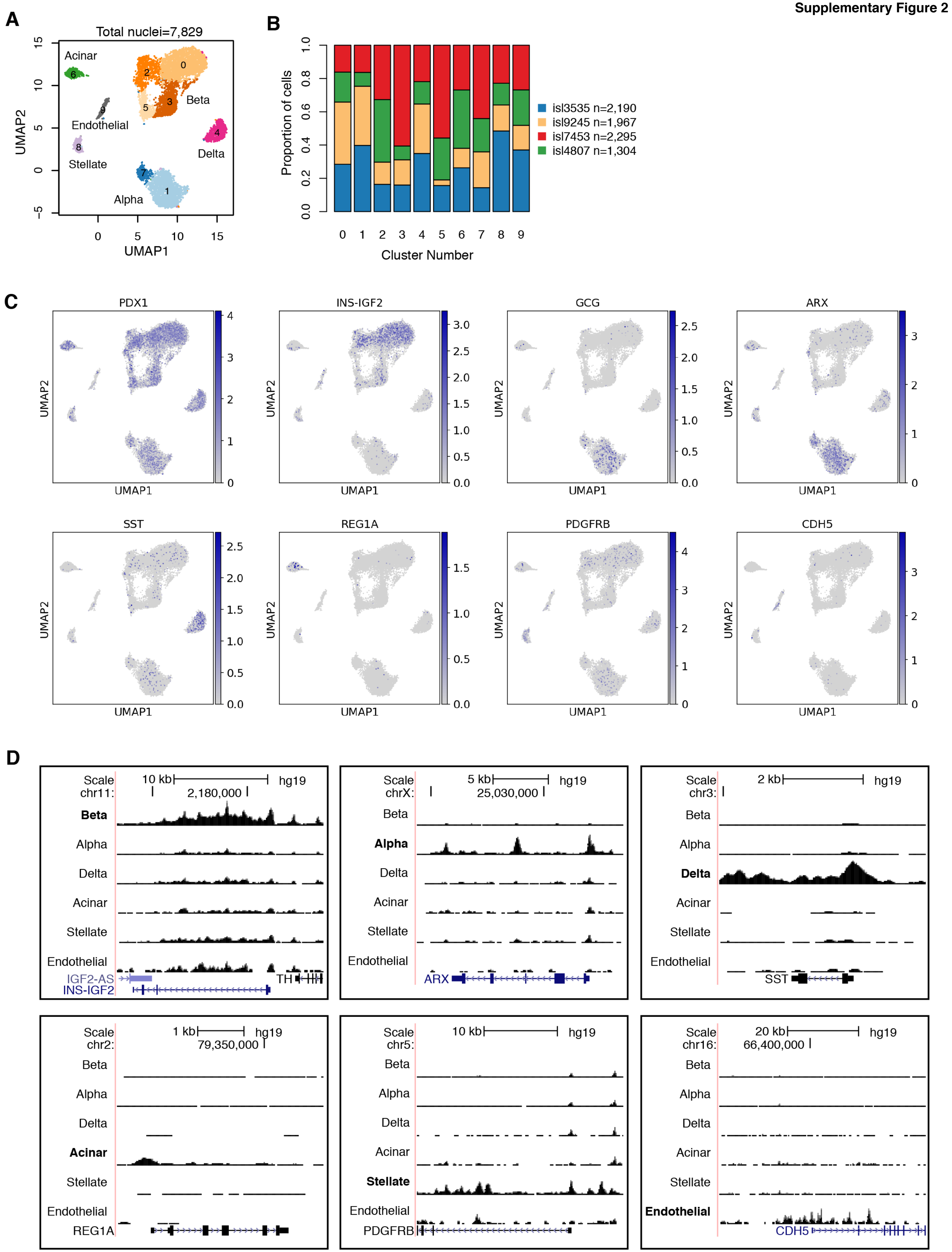
A
Total nuclei=7,829
Acinar
Beta
Endothelial
Delta
Stellate
Alpha
UMAP1
UMAP2
B
Proportion of cells
Cluster Number
isl3535 n=2,190
isl9245 n=1,967
isl7453 n=2,295
isl4807 n=1,304
C
PDX1
INS-IGF2
GCG
ARX
SST
REG1A
PDGFRB
CDH5
D
Scale
hg19
chr11:
2,180,000
chrX:
25,030,000
chr3:
chr2:
79,350,000
chr5:
chr16:
66,400,000
Beta
Alpha
Delta
Acinar
Stellate
Endothelial
IGF2-AS
INS-IGF2
TH
ARX
SST
REG1A
PDGFRB
CDH5

Supplementary Figure 3

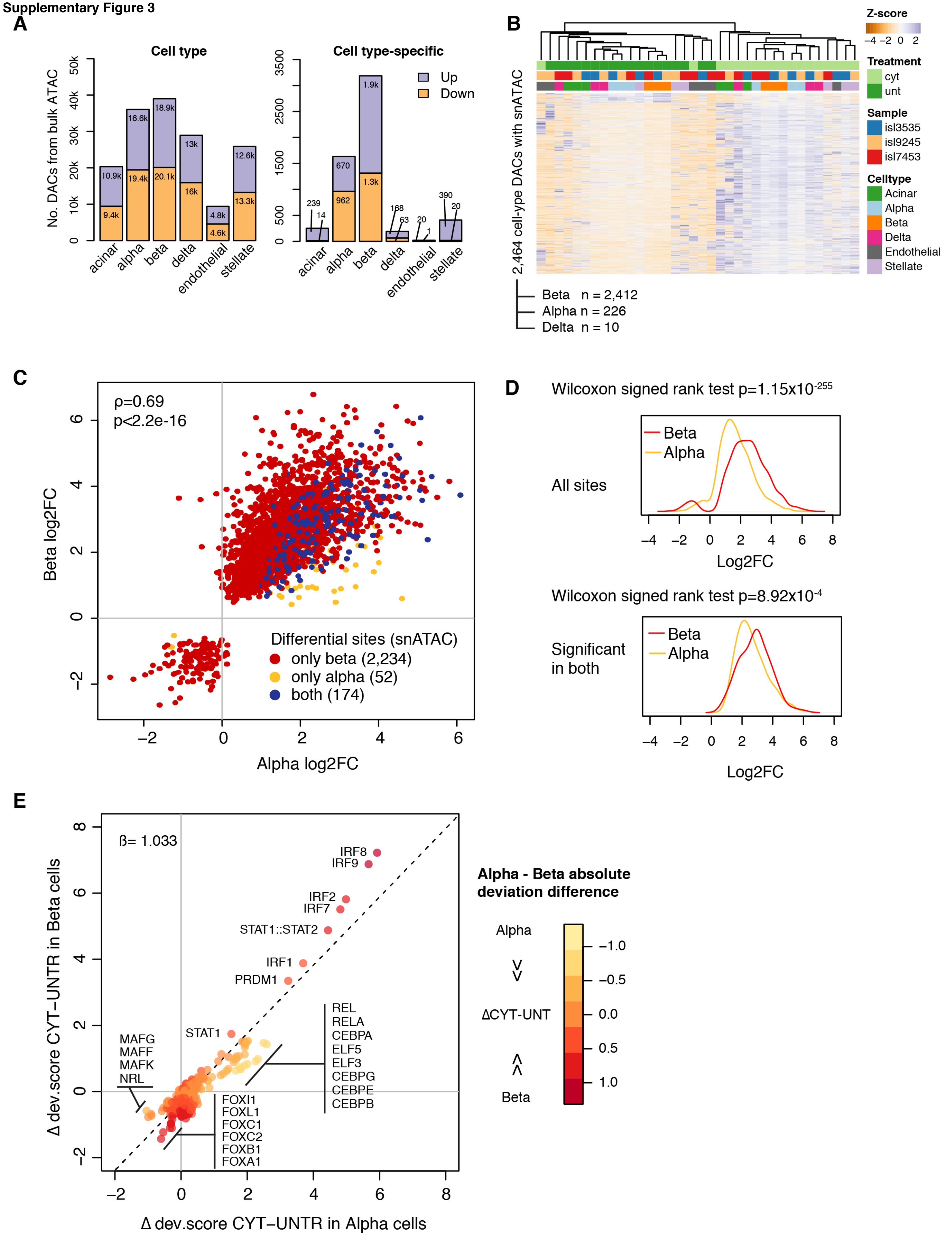

Supplementary Figure 4

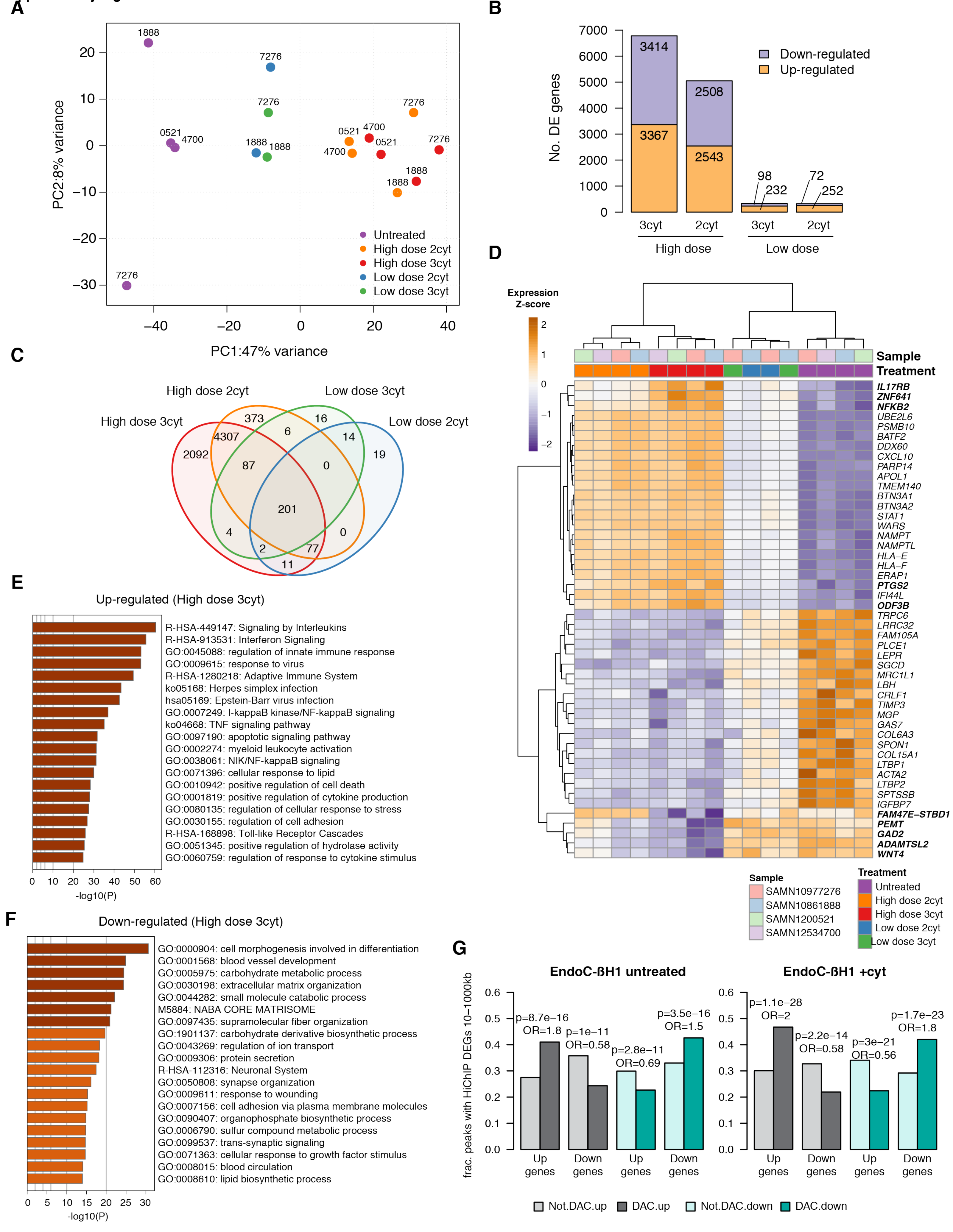

Supplementary Figure 5

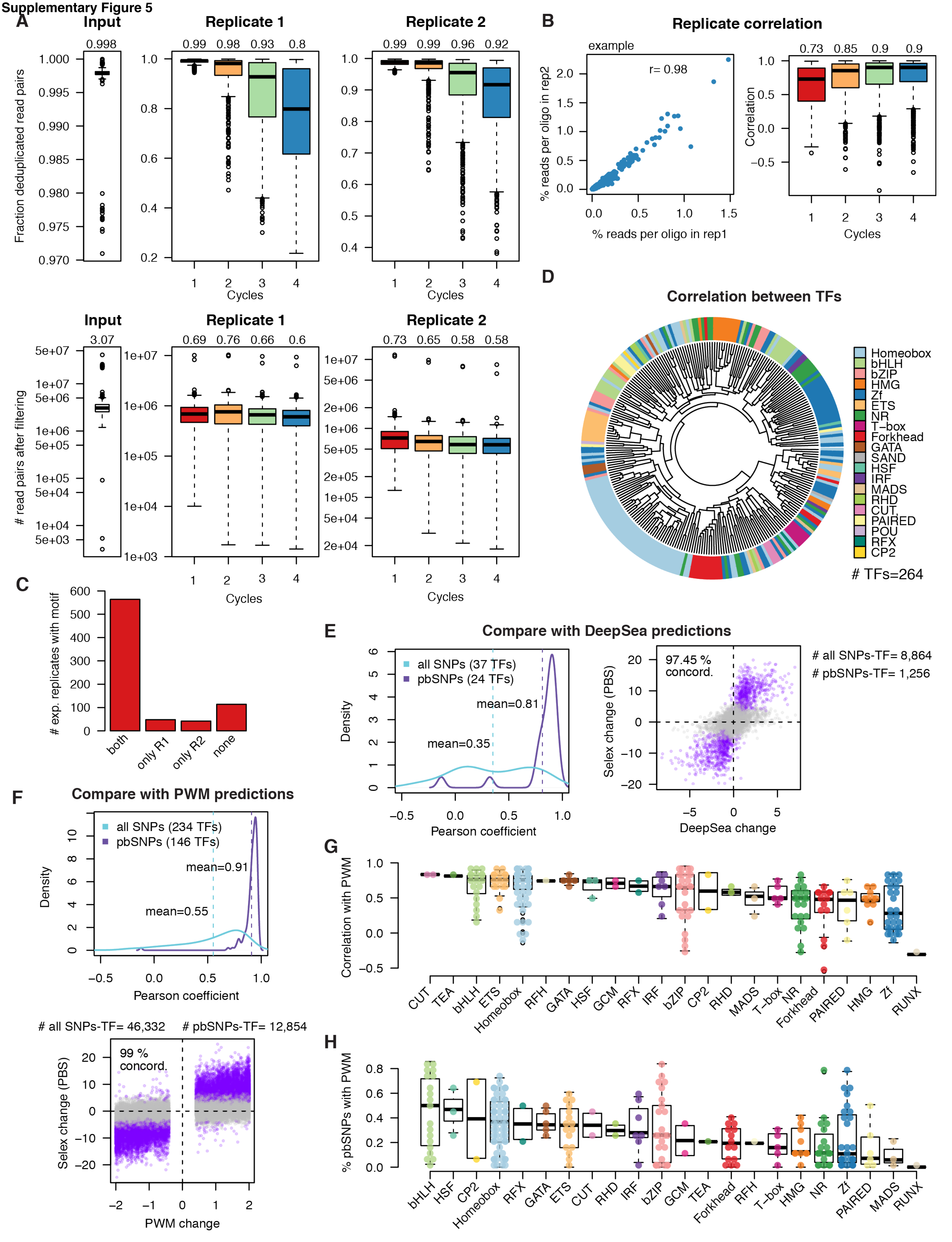

Supplementary Figure 6

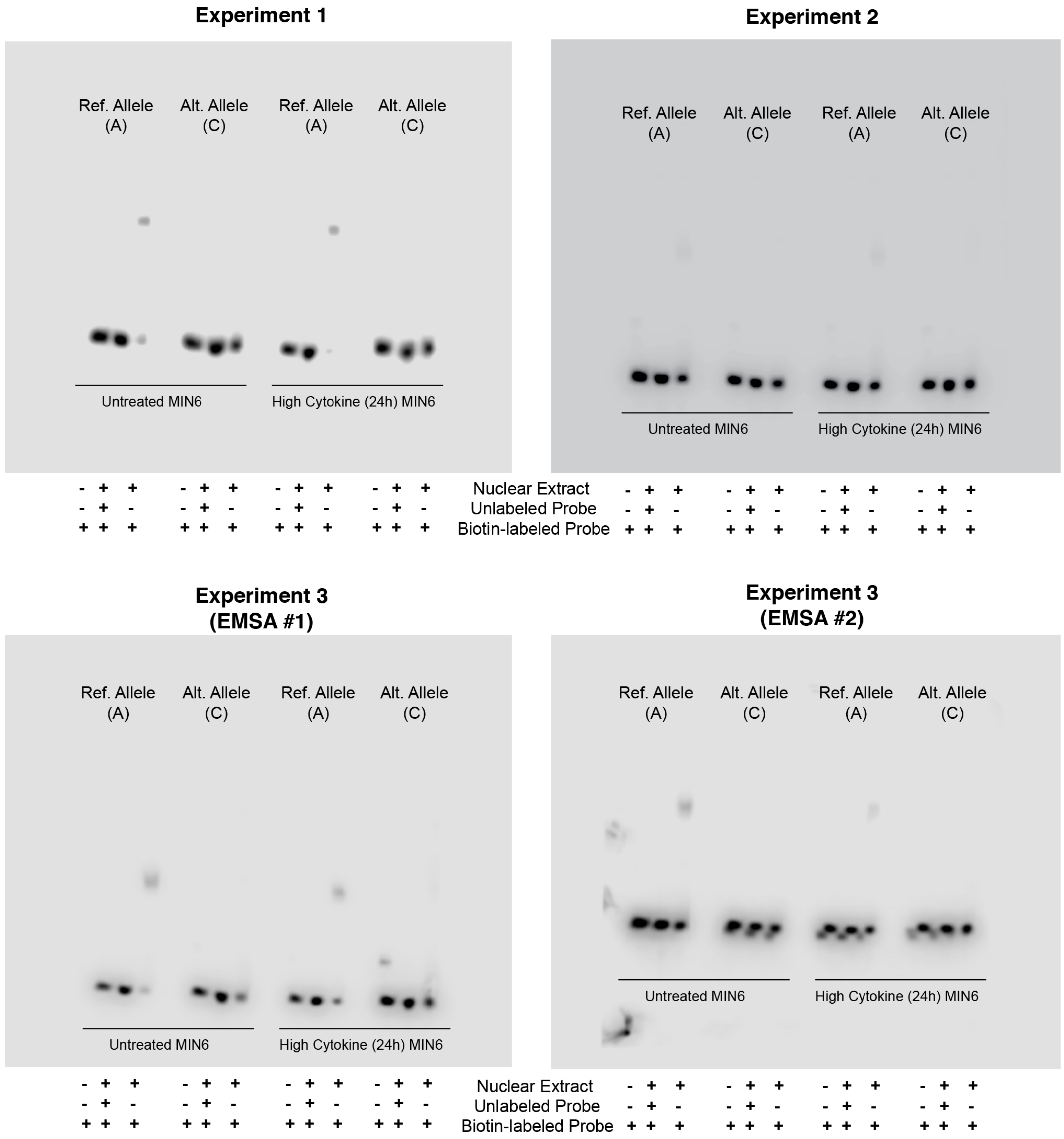
